# Shrinkage Classification for Overlapping Time Series: An interpretable method for mapping stimulus-differentiated evoked response

**DOI:** 10.1101/733279

**Authors:** Peter W. Elliott, Matthew J. Boring, Yuanning Li, R. Mark Richardson, Avniel Singh Ghuman, Max G’Sell

## Abstract

Multivariate time series from neural electrophysiological recordings are a rich source of information about neural processing systems and require appropriate methods for proper analysis. Current methods for mapping brain function in these data using neural decoding aggregate information across space and time in limited ways, rarely incorporating spatial dependence across recording locations. We propose Shrinkage Classification for Overlapping Time Series (SCOTS), a neural decoding method that maps brain function, while accounting for spatio-temporal dependence, through interpretable dimensionality reduction and classification of multivariate neural time series. SCOTS has two components: first, overlapping clustering from sparse semi-nonnegative matrix factorization gives a data-driven aggregation of neural information across space; second, wavelet-transformed nearest shrunken centroids with sparse group lasso performs multi-class classification with selection of informative clusters and time intervals. We demonstrate use of SCOTS by applying it to human intracranial electrophysiological and MEG data collected while participants viewed visual stimuli from a range of categories. The method reveals the dynamic activation of brain regions with sensitivity to different object categories, giving insight into spatio-temporal contributions of these neural processing systems.

## 1. Introduction

Neural electrophysiological recordings provide multivariate time series that reveal dynamic neural processing during experimental tasks. Proper analysis of these time series is challenging due to the high spatial dimensionality and high degree of complexity in the relationships within the recorded activity, with dependence across multiple time scales and spatial scales. Moreover, there are many possible modes of information representation in the brain, including mean-shifts, oscillatory signals, and shifts in statistical dependence between spatial areas.

One important technique for understanding neural processing systems is neural decoding, the process of predicting experimental conditions based on recorded neural activity. If recorded neural activity is sufficient for distinguishing between experimental conditions it implies that the neural signal carries information relevant to the varying conditions. With well-designed experimental conditions, researchers can control what relevant information is captured in the decoding process. However, it is also important to be able to interpret a classifier that successfully discriminates amongst experimental conditions (Yamashita et al., 2008; Rasmussen et al., 2012; Haufe et al., 2014). A black-box classifier that predicts experimental conditions successfully may confirm the presence of relevant signal in recorded neural activity but gives little additional insight. Decoding may also capture information that is not relevant to cognition (Kriegeskorte and Douglas, 2018). Classifiers that maintain some degree of interpretability are advantageous for understanding how information is encoded in neural activity and lead to broader insight into how neural systems process information (Naselaris et al., 2011).

The type of interpretability needed depends on the analysis task. One important task in understanding neural processing is mapping where and at what latency the brain responds to different visual stimuli. Spatial and temporal localization of neural response reveal organizing principles of how the brain processes information and which areas contribute to different stages of processing. Our goal in this paper is to provide an interpretable classification method for locating stimulus-specific neural responses in neural electrophysiological recordings that addresses the challenges posed by high dimensionality and dependence across space and time.

We apply our proposed method to both intracranial EEG (iEEG) and MEG experiments, with a focus on analysis of single trial local field potentials (LFPs) and event related potentials (ERPs). We demonstrate that our method is appropriate for both recording techniques. iEEG is a tool for neuroscientific research in clinical settings that offers high signal to noise ratio and temporal resolution but presents several unique challenges for analysis. Subjects in iEEG studies typically have electrodes implanted primarily for clinical rather than research purposes, with locations of electrodes varying across patients as needed. Additionally, electrodes are not spread uniformly and are instead in clusters of varying size. MEG provides whole brain coverage at lower spatial resolution, with comparable coverage across all subjects. An overview of electrophysiological recording techniques is given by Buzsáki et al. (2012).

### 1.1. Previous Methods

Previous methods for localizing spatial and temporal responses have relied on searching through recording locations and time bins to find discriminative power between experimental conditions. As examples, Liu et al. (2009) applied a linear support vector machine to time-binned features derived from iEEG ERPs to classify stimuli from individual trials. Ghuman et al. (2014) used a time-binned nearest neighbor classifier on single-channel iEEG ERPs to evaluate sensitivity to face images in the fusiform face area. Hirshorn et al. (2016) evaluated word reading sensitivity in the left midfusiform gyrus using time-binned Gaussian naive Bayes. As an alternative approach, Li et al. (2017) proposed connectivity maps based on canonical correlation analysis to relate information in two neural regions of interest and applied the method to decoding images from fMRI and individual face identity from iEEG wavelet features.

This general search strategy has two primary drawbacks. First, the process of searching carries a risk of overfitting due to multiple testing, and correcting for this risk leads to a considerable reduction in statistical power. Second, this strategy limits the ability to recognize and make use of relationships and interactions across spatial locations as each section of the recording is evaluated only in isolation.

A similar “searchlight” method was proposed by Kriegeskorte et al. (2006) for finding informative regions in fMRI, with spherical regions of voxels independently measured for information distinguishing experimental conditions. A spatio-temporal searchlight method has also been proposed for a similar analysis of scalp EEG and MEG imaging (Su et al., 2012). Drawbacks to searchlight analysis are discussed in Etzel et al. (2013). One concern in particular is the size of the searchlight used. A searchlight that is too small may fail to detect a large pool of voxels that are weakly informative individually but strongly informative collectively. If the searchlight is too large, a small number of strongly informative voxels may bias the measure of many nearby uninformative voxels. This underscores the importance of a data-driven approach to extracting the main modes of variation in neural response instead of uniform search.

### 1.2. Our Proposal

In this paper we propose Shrinkage Classification for Overlapping Time Series (SCOTS), a novel method for dimensionality reduction and interpretable classification that aggregates information from neural activity across space and time in a data driven manner. SCOTS follows two steps. In the first step we generate a low rank representation of the multivariate time series using sparse semi-nonnegative matrix factorization. This factorization finds both overlapping clusters of recording channels and corresponding latent cluster-level time series. In the second step we estimate condition-specific deviations of the latent time series from the overall signal mean using wavelet-transformed nearest shrunken centroids with an additional sparse group lasso penalty. We can validate discovered condition-specific deviations using out-of-sample nearest centroid classification.

Classical nonnegative matrix factorization (NMF) (Lee and Seung, 1999) is a linear dimensionality reduction method that approximates a data matrix **X** as the product of two matrices **W** and **H**, each with nonnegative entries. Rows of the right hand matrix **H** form a basis for rows of **X**, while rows of the left hand matrix **W** serve as basis coefficients or loadings. NMF is appropriate for data that is naturally nonnegative and built from layers of components and therefore has has been applied in a wide variety of contexts, including characterization of high gamma response profiles to speech in iEEG (Hamilton et al., 2018), tumor detection in spectroscopic imaging (Ortega-Martorell et al., 2012; Sajda et al., 2004), community detection (Wang et al., 2011), audio analysis (Févotte et al., 2009), computational biology (Devarajan, 2008), document clustering (Xu et al., 2003), and blind source separation (Virtanen, 2007). Semi-NMF (SNMF) (Ding et al., 2010) relaxes the nonnegativity constraint on the basis matrix. This is appropriate when data is not nonnegative but is still built from layers of components. Penalized versions have also been proposed to promote smoothness in the basis vectors and sparsity in both **W** and **H** (Cichocki et al., 2007; Drakakis et al., 2008; Yokota et al., 2015). A comprehensive overview of extensions is given by Wang and Zhang (2013).

In our method, SNMF represents time series observed from recording channels as nonnegative mixtures of unconstrained latent time series. We impose additional sparsity on these non-negative mixtures via an *L*_1_ penalty, so that each recording channel is built up from a small number of latent time series, and each latent time series appears in a small number of recording channels. We also project the latent time series onto a truncated wavelet basis to account for dependence across time. Unlike with searchlight methods, the estimated latent time series aggregate neural activity in a way that preserves observed variation rather than in pre-defined windows. Sparsity in this low dimensional representation also helps with spatial mapping of neural responses to stimuli.

Nearest shrunken centroids (NSC) was first proposed by Tibshirani et al. (2003) for interpretable classification, associating genes with diseases. The goal of NSC is to perform multi-way classification with class-specific variable selection. The variables to be selected in our method are time points in a specific latent time series. This poses a challenge because separate time points are not independent – we should select time intervals rather than scattered time points. We address this problem by performing NSC on the coefficients in a wavelet basis for the latent time series. As a result, variable selection picks a sparse set of wavelet basis elements, which correspond to a curve on a compact time interval as desired. Wavelet shrinkage and sparsity has been established as a successful tool for adaptive curve estimation (Donoho and Johnstone, 1994; Zhao et al., 2012). We additionally use a group lasso penalty so that entire non-discriminative clusters can be eliminated.

We emphasize the goal of interpretability in designing this method. Being able to attribute classification success to certain spatial regions and temporal latencies helps us learn more about underlying neural processing than using a black box classifier with superior predictive performance. Sparsity plays a critical role in the interpretability of our method, both in the dimensionality reduction provided by SNMF and the class-specific variable selection of wavelet nearest shrunken centroids.

### 1.3. Application to a Visual Category Localizer Experiment

As an illustration of the proposed method, we applied SCOTS to human subject iEEG and MEG data from a visual category localizer experiment. In the experiment, subjects were shown a series of images, each falling into one of several categories, and they were asked to indicate if the same image was repeated consecutively. The experiment focused on neural activity in the ventral visual stream, an area that is believed to be particularly important for object recognition (Kravitz et al., 2013). Characteristic ERPs of responses in the ventral visual stream to various visual stimuli have been described by Allison et al. (1994); Nobre et al. (1994); Allison et al. (1999); McCarthy et al. (1999). We are interested in measuring category-specific activity in this experiment as it is indicative of an area’s involvement in processing a particular visual concept. Spatial and temporal localization are particularly important as they can reveal at what stage of visual processing a certain concept is distinguished and improve understanding of the dynamics of visual processing as a whole.

We applied SCOTS to visually-evoked LFPs from across each of nine iEEG subjects and three MEG subjects. The number of recording channels per iEEG subject ranged from 22 to 144 with placement depending on a mix of clinical and research goals. Analysis for MEG subjects was performed in source space in the left hemisphere. Results derived from both imaging modalities provided interpretable dynamic neural activity that was centered in cortical regions previously implicated with visual or language processing.

## 2. Methods

### 2.1. Overview

We propose SCOTS, a method for spatio-temporal localization of condition-specific mean shifts in ERPs in response to stimuli. The method has two stages: in Section 2.2, sparse SNMF gives soft clustering of recording channels and separation of latent source signals; in Section 2.3, a grouped version of nearest shrunken centroids is applied in a wavelet transformed space for selection of cluster-category pairs and corresponding time regions with distinct behavior. These two stages are illustrated in Figure 2.1. To validate discovered spatio-temporal category sensitivity, we classify held-out trials using fitted centroids (Section 2.4).

**Figure 1:**
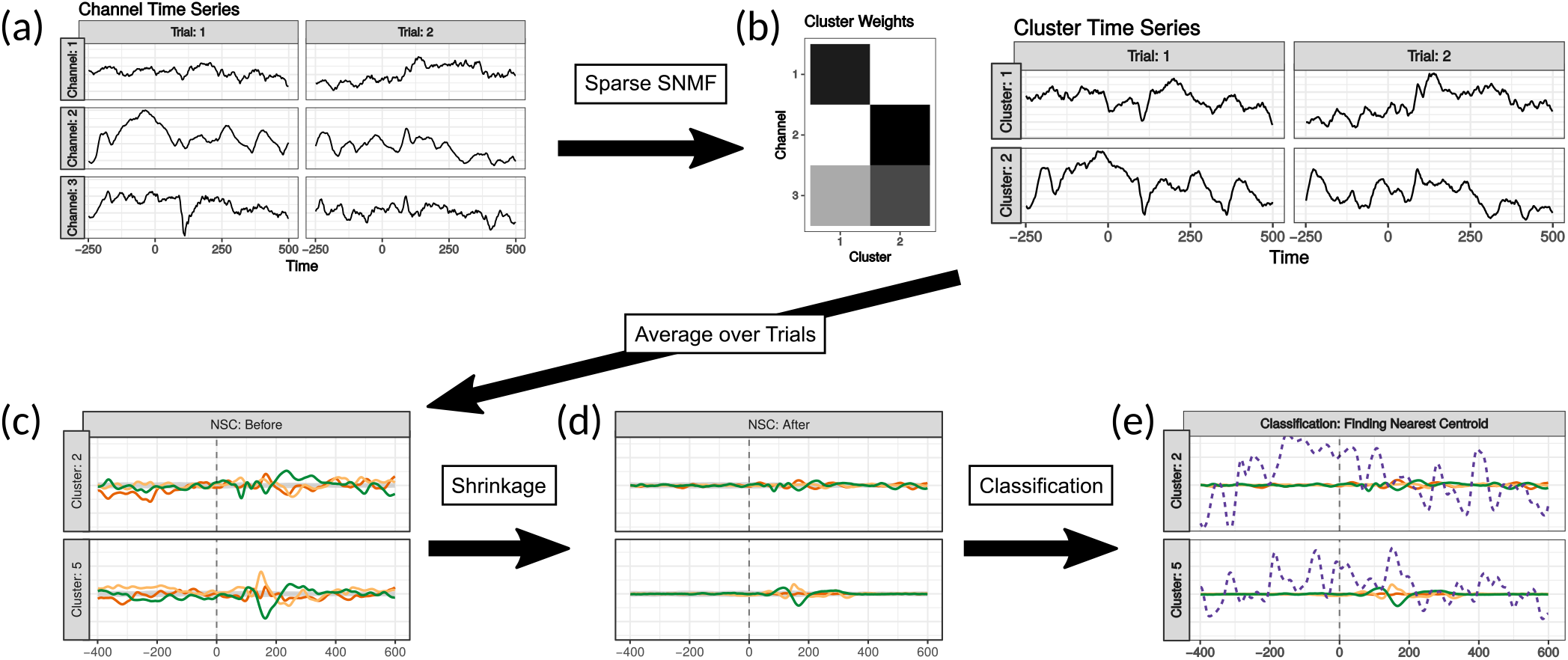
SCOTS Overview: (a) we observe *p* recording locations for *n* trials and *T* time points per trial. (b) We apply sparse SNMF to get *k* overlapping clusters of recording locations, with associated weights and cluster time series for each trial (Section 2.2). (c) We average cluster time series over trials, then (d) apply NSC to get centroid time series for each cluster and trial category (Section 2.3). NSC uses shrinkage to find clusters and time ranges with category sensitivity. The gray confidence band for the overall mean across categories is used to decide the amount of shrinkage – all category centroids are within the confidence band before the patient is shown an image (Section 2.5.4). (e) New observations, e.g. the purple dashed time course, are classified by finding the nearest centroid.

**Figure 2:**
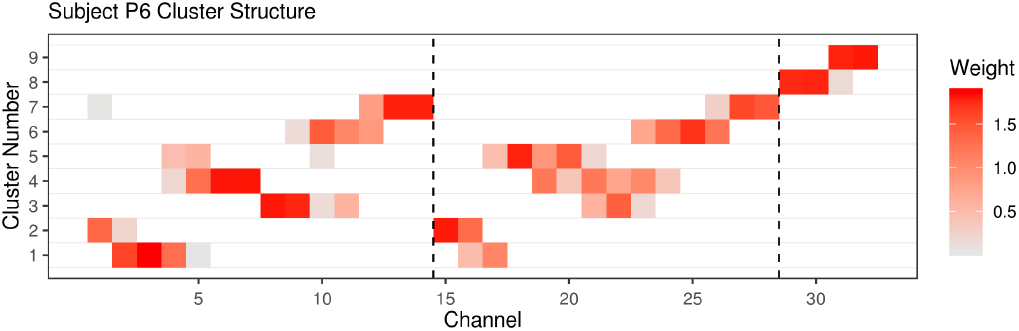
P6 Cluster Structure: This plot shows basis coefficients mapping recording channels to clusters for subject P6. The first 28 recording channels are located in the left basal temporal area, and the last four are located in the left temporal pole. Channels 1-14 and 15-28 run in parallel on high density strips running from lateral to medial on ventral temporal cortex (see Figure 5). This structure is reflected in the cluster weights.

### 2.2. Sparse Semi-Nonnegative Wavelet-Projected Matrix Factorization

The first stage of SCOTS uses a sparse adaptation of SNMF with a truncated wavelet basis. SNMF builds an approximation **X** ≈ **WH**, where **X** is a data matrix, **H** is an unconstrained basis matrix, and **W** is a nonnegative coefficient matrix. The choice of nonnegativity in **W** reflects an interpretation of observations in **X** as being built from additive layers of basis vectors in **H**, where the basis vectors are latent time series.

In our application we observe a matrix **X**_*i*_ ∈ ℝ^*p*×*T*^, where *p* is the number of electrodes and *T* is the number of time points, for each of *i* = 1,…, *n* experimental trials, and we decompose each observed matrix in the wavelet domain: **X**_*i*_Φ ≈ **WH**_*i*_Φ, with Φ a pre-determined truncated wavelet basis matrix. Our version of SNMF is defined by the following optimization problem, with **F**_*i*_ = **H**_*i*_Φ consisting of wavelet basis coefficients:

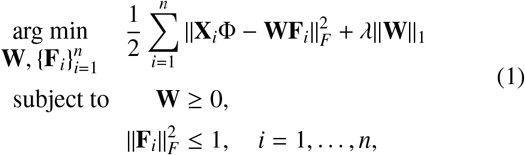

where *K* is the number of latent time series, **W** ∈ ℝ^*p*×*K*^, **F**_*i*_ ∈ ℝ^*K*×*T*^, and *λ* is a penalty weight. We can recover a time-domain decomposition of **X**_*i*_ by setting **H**_*i*_ = **F**_*i*_Φ^*T*^.

The coefficients **W**, which are fixed across all trials, can be interpreted as determining a set of clusters of variables with shared behavior. Sparsity in the coefficients is important for this interpretation, as such clusters may overlap but should be relatively local in space. This is especially the case because modeling **W** as fixed across time and categories implies that clusters should mostly reflect spatial propagation of electrophysiological neural activity. The nonnegativity constraint imposes some sparsity but is generally not sufficient for this interpretation. We therefore impose additional sparsity is by adding an *L*_1_ penalty to **W** in (1).

We perform SNMF in the wavelet domain to account for time dependence in observations within each trial. A wavelet basis represents a univariate time series as a linear combination of basis vectors, each associated with a time shift and a time scale. Wavelet bases are popular in signal processing for their ability to represent curves with sometimes sharp changes using relatively few basis vectors. After transforming back into the time domain, the use of a truncated wavelet basis results in smooth latent time series **H**_*i*_ that can also have sharp jumps, as have been observed in characteristic ERPs for visual stimuli (Allison et al., 1994; Nobre et al., 1994; Allison et al., 1999; McCarthy et al., 1999). The truncated wavelet basis projection also offers computational benefits, as the number of wavelet coefficients used is much smaller than the number of time points *T*.

### 2.3. Wavelet Nearest Shrunken Centroids

After performing dimensionality reduction, SCOTS finds localized category-specific neural responses by applying a modification of nearest shrunken centroids to the latent time series **H**_*i*_. This finds spatio-temporal regions where a centroid for a category can be distinguished from the average across all categories. Nearest shrunken centroids was originally proposed for diagnosing medical conditions using gene expression data. In that context, sparsity in gene-diagnosis associations was used to simplify the interpretation of a diagnosis. The basic strategy is to shrink within-category means for each covariate to the overall mean using soft-thresholding adjusted for covariate-specific variance. This strategy is unsatisfactory when applied to multivariate time series, since each latent time series has natural dependence across time that would not be accounted for. We should also expect cluster-wise sparsity. That is, if the response for a category deviates from the overall average, that deviation should only appear in a small number of latent time series.

We address time-dependence by projecting the differences between each category centroid and the overall mean onto a wavelet basis for each latent source. Switching to the wavelet domain allows us to leverage the sparsity encouraged by nearest shrunken centroids to get curves for the differences between centroids and the overall mean that respect time dependence. Use of wavelets also gives additional control over time localization. As we use a truncated basis, i.e. set all basis coefficients to zero for the finest time scales, we get an initial smoothing of the centroids across time. We additionally can weight thresholding differently across wavelet time scales, which allows us to set a stricter standard for finding differences in neural response on time scales that are biologically less likely.

We address the need for cluster-wise sparsity by adding a group lasso penalty to the differences between the category centroids and the overall mean. We group together all time points for each cluster-category pairing. This encourages entire latent time series to show no difference between a given category centroid and the overall mean, thus restricting estimates of differentiated neural response to a small number of recorded locations.

Our modified version of nearest shrunken centroids can be written as follows. Suppose *g*(*i*) is the category label for trial *i*, *G* is the number of categories, and **W** and **H**_*i*_ are the factors estimated using sparse SNMF. Further, let Φ be a truncated wavelet basis matrix and *w* be the weights determined by the wavelet scales. We use **A**_*k*,:_ to denote row *k* of matrix **A**. We solve the following optimization problem:

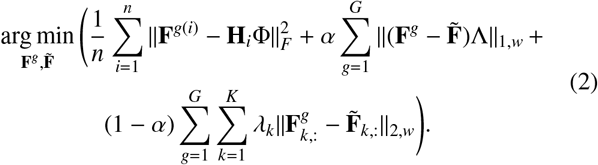

Here Λ is a diagonal matrix with entries *λ*_1_,…, *λ*_*K*_ as cluster-specific penalty parameters controlling the overall penalization. An additional parameter *α* ∈ (0, 1) controls the trade-off between individual and group sparsity. The matrices **F**^*g*^ and 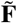 represent the wavelet coefficients for the category *g* centroid and the overall mean respectively. To get category centroids back in the time domain we simply invert the wavelet transformation: **H**^*g*^ = **F**^*g*^Φ^*T*^.

### 2.4. Classifying New Observations

After finding category centroids in latent time series space, we can classify held-out trials to validate apparent differentiated responses. Suppose that we have estimated **W** and centroids **H**^*g*^ on a training set and want to classify a new trial **X**^+^ not used for training. We estimate the latent time series for the new trial by projection using **W**. The standard least squares estimate is **H**^+^ = (**W**^*T*^ **W**)^−1^**W**^*T*^**X**^+^. The distance to each estimated centroid is *d*(**H**^+^, **H**^*g*^) = ║**H**^+^ − **H**^*g*^║_*F*_. The new observation is then classified according to the minimum distance.

### 2.5. Visual Category Localizer Experiment

#### 2.5.1. Experimental Setup

We applied SCOTS to a visual category localizer experiment where human subjects were shown a sequence of images, each from one of several categories. Neural activity in nine subjects was recorded using iEEG while the images were shown. Results from this experiment and details regarding data collection and pre-processing were previously published in Ghuman et al. (2014). We also analyzed three subjects for whom neural activity was recorded using MEG.

For iEEG subjects, the number of recording channels used ranged from 22 to 144. The area of greatest interest was the ventral temporal cortex. For subjects with fewer than 20 ventral temporal recording channels we used all recording channels from the experiment. Details for the recordings are shown in Table 2.5.1. Two of the subjects, P2 and P4, participated in two sessions each, labeled (a) and (b) respectively. For each category there were 30 images. Each image was shown a baseline of twice, with a random of chance of being repeated on the following trial. This repeat probability was either 1/3 or 1/6 depending on the subject. The sets of images of faces and bodies were each 50% male.

**Table 1:**
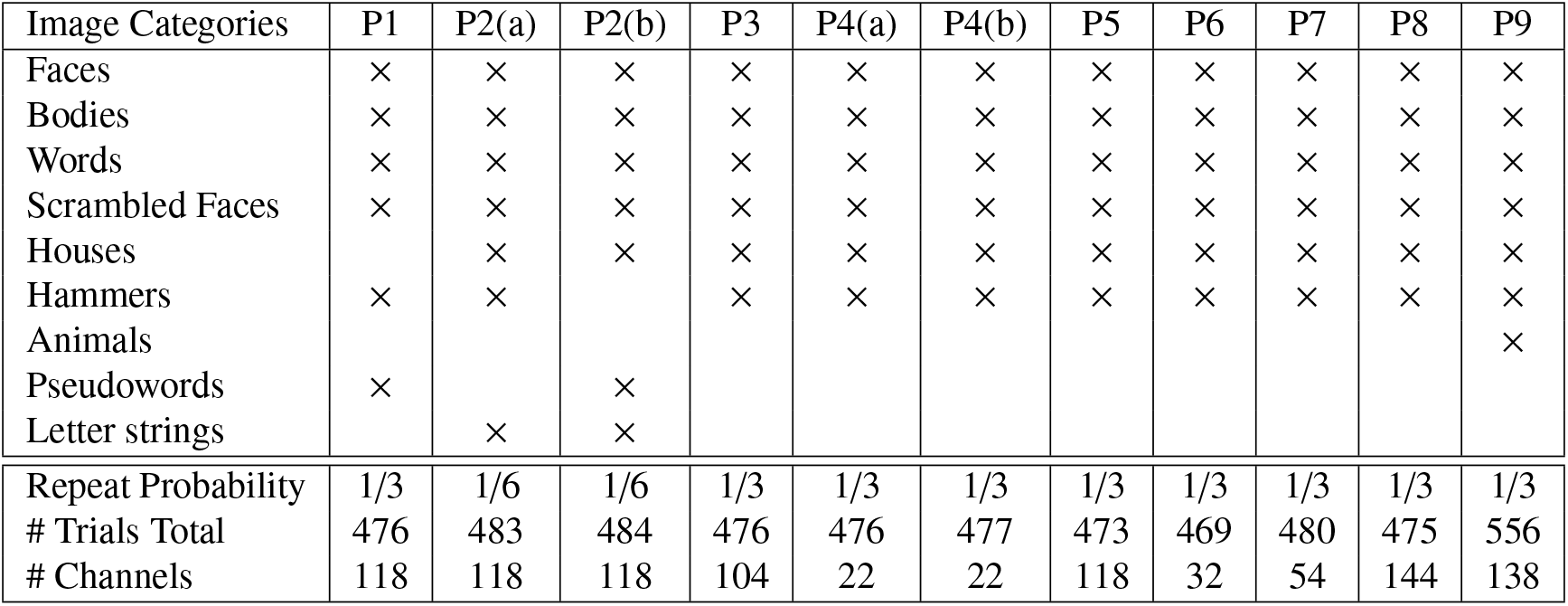
Protocol Details: Each subject was shown images from several categories. Here we indicate the categories presented to each iEEG subject, P1-P9, with (a) and (b) indicating repeats of the experiment with the same subject. The listed repeat probability is the proportion of trials for which the same image was presented twice in a row, to which the subject responded with a button-press. The number of channels is the number of recording locations we analyzed from each subject.

**Table 2:** NSC-Selected Category Sensitivity: For each subject, we use NSC to select clusters with category-specific neural activity. Here we list for each subject the categories with sensitive clusters detected by NSC with false discovery rate (FDR) control. Bold indicates categories that were also selected with family-wise error rate (FWER) control.

| Subj. | Categories |
| --- | --- |
| P1 | Faces, Scrambled Faces |
| P2(a) | × |
| P2(b) | Houses, Scrambled Faces |
| P3 | × |
| P4(a) | **Faces, Words, Scrambled Faces, Houses, Bodies, Hammers** |
| P4(b) | **Faces, Words, Scrambled Faces, Houses** |
| P5 | **Words, Scrambled Faces** |
| P6 | **Faces, Words, Scrambled Faces, Bodies, Hammers** |
| P7 | **Words** |
| P8 | **Faces, Words, Scrambled Faces, Houses, Bodies** |
| P9 | **Faces, Words, Scrambled Faces, Houses, Hammers** |

In each trial in each session, the image was presented for 900 ms with 900 ms inter-trial interval during which a fixation cross was presented at the center of the screen (approximately 10° × 10° of visual angle). Participants were instructed to press a button on a button box when an image was repeated (1-back).

For MEG subjects, analysis was performed in source space in the left hemisphere, with the number of vertices ranging from 3,606 to 3,853. Subjects were shown images from four categories: words, houses, hammers, and false fonts. For each category there were 30 images. Each image was shown a baseline of three times with a 1/6 chance of being repeated on the following trial, for a total of 420 trials. The images of faces were 50% male. In each trial the image was presented for 300 ms with a 1.5 s inter-trial interval.

Each of the iEEG subjects were patients with pharmacologically intractable epilepsy that had intracranial electrodes implanted for the localization of epileptic foci. Ages of the iEEG subjects ranged from 19 to 56 with five females. The MEG subjects were healthy control participants. Ages of the MEG subjects ranged from 19 to 27 with one female. All patients and subjects provided written informed consent to experimental protocols approved by the University of Pittsburgh’s Institutional Review Board.

#### 2.5.2. MEG and iEEG Data Pre-Processing

Local field potentials were recorded from iEEG electrodes via a GrapeVine Neural Interface (Ripple, LLC) with 1 kHz sampling rate. Data was subsequently band pass filtered from 0.1-115 Hz using fourth order butter-worth filters, and notch filtered at 60/120/180 Hz using the FieldTrip toolbox (Oostenveld et al., 2011). Data was then epoched from 500 ms pre to 1000 ms post-stimulus presentation. iEEG electrodes were localized to individual subject anatomy via post operative MRIs or CT scans using the brainstorm MATLAB toolbox (Tadel et al., 2011). Surface electrodes were projected to the nearest vertex on the pre-operative MRI to correct for brain shift (Hermes et al., 2010). Trials with outlying neural activity were removed (see Appendix B).

MEG data were collected on an Elekta Neuromag Vectorview system (Elekta Oy, Helsinki, Finland) with 204 gradiometers and 102 magnetometers arranged in orthogonal triplets. Data were collected at 1000 kHz, then subsquently pre-processed with Signal-Space Projection operators tailored to eliminate environmental artifacts based on empty room data (Tesche et al., 1995; Uusitalo and Ilmoniemi, 1997). Data were then band-pass filtered from 1-50 Hz (5 Hz low-pass transition band, 3-sample high-pass transition band and 8917 sample filter length) then downsampled to 250 Hz using the MNE-C toolbox (Gramfort et al., 2014). Finally, data were pre-processed via temporal source space separation (10-second buffer length, 0.98 correlation limit) (Taulu and Hari, 2009; Taulu and Simola, 2006) using Elekta MaxFilter software. Data were then epoched from −500 to 1000 ms around stimulus presentations.

#### 2.5.3. Training and Testing

For each subject, experimental trials were split into training and testing sets. The training sets consisted of 36 randomly chosen trials from each image category. The testing sets consisted of all other trials. Sparse SNMF was applied only to the training set for each subject. Trials in the test set for each subject were projected onto the corresponding estimated weight matrix **W**, as in Section 2.4.

#### 2.5.4. Choosing Tuning Parameters

Applying sparse SNMF required choosing a number of clusters and a sparsity penalty parameter for each subject. For each iEEG subject we solved the sparse SNMF problem for a grid of *k* and *λ* values. The best parameters were chosen to balance two criteria: (i) increase in reconstruction loss when decreasing *k* and/or increasing *λ*, and (ii) stability of cluster estimates under bootstrapping of training trials. The first criterion is analogous to the scree plot elbow rule for choosing dimension in PCA. To reduce computational burden for MEG subjects, a common *k* was chosen based on singular values of the data matrices.

Applying wavelet nearest shrunken centroids required choosing penalty mixture *α* and cluster sparsity parameters *λ*_1_,…, *λ*_*K*_. The cluster sparsity parameters were chosen so that category centroids fell within a 90% confidence band for the overall mean in the pre-stimulus LFP. The confidence band for the overall mean was estimated using a bootstrap sample of the training set, without stratifying by category. The penalty mixture *α* was set to 0.9, with results robust to the choice of the parameter. We could alternatively select *λ*_1_,…, *λ*_*K*_ via cross validation within the training set, but we prefer using the pre-stimulus as a control because it tends to give sparser, and therefore more interpretable, models.

#### 2.5.5. Single Category d′

Although we perform multi-way classification, only a subset of categories may be possible to classify. This is because the recorded areas of the brain are likely only involved in processing particular types of visual stimuli; our ability to accurately classify experimental trials is indicative of which stimuli the area is involved in processing. Selective classification accuracy is therefore of scientific interest rather than a statistical failing.

We account for category-specific classifiability by measuring sensitivity, the proportion of trials from a given category that are accurately labeled, and false positive rate, the proportion of trials from all other categories that are labeled as being from the given category, separately for each category. For each category we assess sensitivity and false positive rate using a single index, *d*′, related to the Gaussian inverse CDF Φ^−1^ by

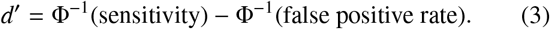

We use *d*′ as a measure of separation because it tends to be reasonably insensitive to classification cutoff; indeed, under certain assumptions it is invariant to cutoff (Macmillan and Creelman, 2004). This allows us to make reasonable comparisons even between classifiers tuned for different trade-offs between sensitivity and false positive rate.

##### 2.5.6. Single Cluster Classification

As another test of localized information content, we assessed modified classifiers that were restricted to using only one cluster at a time. These modified classifiers differ only in the final stage – for each cluster *k* we can calculate 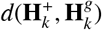, the distance from the projection for the new trial to the shrunken centroid for category *g* restricted to cluster *k*. This helps to summarize the discriminating information present in each cluster.

##### 2.5.7. Permutation Tests for Baseline Category Deviation and Classification Performance

We used permutation tests to select clusters that showed behavior unique to a particular category as estimated by wavelet nearest shrunken centroids. We assessed the deviation from consensus behavior in each cluster *k* and category *g* by finding the maximum ratio *ρ*_*k*,*g*_ of the category centroid’s deviation from the overall mean across time to the time-varying bootstrap standard error of the overall mean:

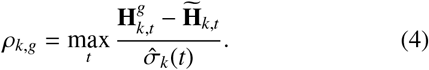

This statistic both accounts for differing baseline variation and the intuition that behavior of interest may be short in duration. Deviation size was assessed by refitting wavelet nearest shrunken centroids for 200 random shuffles of the training labels.

We used two selection schemes: the first controlling familywise error (FWER) across clusters and the second controlling false discovery rate (FDR) across clusters. For each randomly trained model, we calculated the overall maximum deviation ratio *ρ*_max_ = max_*k*,*g*_ *ρ*_*k*,*g*_ and the cluster-wise maximum deviation ratios *ρ*_*k*,max_ = max_*g*_ *ρ*_*k*,*g*_. The overall maximum deviation ratios form a distribution for the largest observed deviation under a global null hypothesis that the true deviation in the mean is zero for all clusters and categories. We then select cluster-category pairs from the non-permuted results with a deviation ratio exceeding the 95^*th*^ percentile of the permutation distribution to get FWER control. The cluster-wise maximum deviation ratios are used to test each cluster separately. We can calculate *p*-values from the permutation distribution for each cluster and use the Benjamini-Hochberg procedure (Benjamini and Hochberg, 1995) to get cluster selections with FDR control.

We also used permutation tests to assess all-cluster classification performance. These permutation tests are used to account for the bias that results from picking the best classification performance across all categories. We repeated the permutation test for single-cluster classification, but for each permutation we used the maximum *d*′ across all cluster-category pairs, restricting to cases where sensitivity was at least 10%. This restriction removed cases where *d*′ was infinite due to lack of false positives but classification was not meaningfully successful because of the very small sensitivity. In both cases permutation tests were based on refitting wavelet nearest shrunken centroids under 200 random shuffles of category labels.

One caveat for the permutation test we used is that the classification performances for different categories are not independent within each subject. In fact, sensitivity to one category will tend to shrink the false positive rate for other categories, resulting in positive correlation across categories. The permutation test is for the global null hypothesis that none of the categories have sensitivity and does not establish the distribution of the *k*^*th*^ best *d*′ under the null hypothesis that there are *k* − 1 truly sensitive categories for *k* ≠ 1. As a result, comparison to permutation test thresholds is only a heuristic assessment for lower ranked categories. This is a generic problem for all permutation tests.

##### 2.5.8. Model Comparisons

We compared our method’s predictive performance on iEEG experiments to two similar procedures. The first procedure is a version of SCOTS with tuning parameters for wavelet nearest shrunken centroids chosen by cross validation (see Section 2.5.4). The second procedure builds a prototype for each class by taking the within-class mean, using neither dimensionality reduction nor centroid shrinkage. For MEG, the cross validated version of SCOTS is replaced with a shrinkage-free version due to computational constraints. The area under the curve (AUC) for each category is calculated by adjusting a classification threshold based on the difference in the distance from the observation to the category centroid to the minimum such distance across all categories.

## 3. Results

### 3.1. Overlapping clusters in subjects

We applied sparse SNMF, described in Section 2.2, to the training set for each subject to find overlapping clusters of recording channels with shared behavior and cluster-level time series for each experimental trial. The cluster structure for one of the iEEG subjects, P6, is shown in Figure 2. These recording channels are split between the left basal temporal area and the left temporal pole. The number of clusters and sparsity penalty were chosen as in Section 2.5.4. This resulted in 19 clusters for 138 channels in P9, 10 clusters for 144 channels in P8, 11 clusters for 54 channels in P7, 9 clusters for 32 channels in P6, 20 clusters for 118 channels in P5, 9 clusters for 22 channels in P4, 17 clusters for 104 channels in P3, 20 clusters for 118 channels in P2, and 20 clusters for 118 channels in P1. All MEG subjects had 40 clusters.

The clusters above came from applying sparse SNMF to time series that included both pre-stimulus and post-stimulus time segments and used a shared decomposition across all categories. We also considered separately modeling the cluster structure for each stimulus category and shorter time segments but found that the results were consistent across category and time. This suggests that the cluster structure was driven by fixed spatial relationships rather than as a response to the experimental stimuli.

### 3.2. Localizing effects

After applying sparse SNMF to obtain cluster-level time series for each trial, we applied wavelet nearest shrunken centroids, described in Section 2.3. This method yields category-specific centroids describing average behavior in each cluster across all trials for the respective category. The deviations of the estimated centroids for P6 from the means from selected categories across all categories are shown as an example in Figure 3. The estimated category sensitivity for each subject based on NSC is shown in Table 3.2. These selected locations, chosen as described in Section 2.5.7, are candidates for having category sensitivity.

**Table 3:** Single-Cluster Classification Results for NSC-Selected Sensitivity: For each subject and each cluster selected by NSC with either FWER or FDR control, we test whether each cluster can be used to independently classify trials on its own (Section 2.5.6). We evaluate success using a permutation test (Section 2.5.7), comparing cluster-specific *d*′ to the best *d*′ when trial labels are shuffled, excluding cases with sensitivity under 10%. Categories with significant single-cluster classification are listed, with bold indicating that the cluster was originally selected by NSC with FWER control. We also report proportions of single-cluster classification significance among clusters selected by NSC, separated by whether the NSC selection was based on FDR or FWER control, showing 82% overall verification after FWER-controlled selection and 68% verification among clusters selected with FDR-control but not FWER-control.

| Subj. | NSC-Selected Categories with Successful Classification | Prop. of FWER | Prop. of FDR |
| --- | --- | --- | --- |
| P1 | Faces, Scram. Faces | 0/0 | 2/2 |
| P2(b) | × | 0/0 | 0/2 |
| P4(a) | **Faces, Words, Scram. Faces, Houses, Bodies, Hammers** | 13/14 | 18/22 |
| P4(b) | **Faces, Words, Scram. Faces** | 5/7 | 7/9 |
| P5 | **Words** | 1/1 | 1/4 |
| P6 | **Faces, Words, Scram. Faces** | 6/7 | 8/12 |
| P7 | × | 0/1 | 0/2 |
| P8 | **Faces, Words, Scram. Faces, Houses, Bodies** | 6/6 | 7/9 |
| P9 | **Faces, Words, Scram. Faces, Houses, Hammers** | 5/8 | 8/15 |

**Figure 3:**
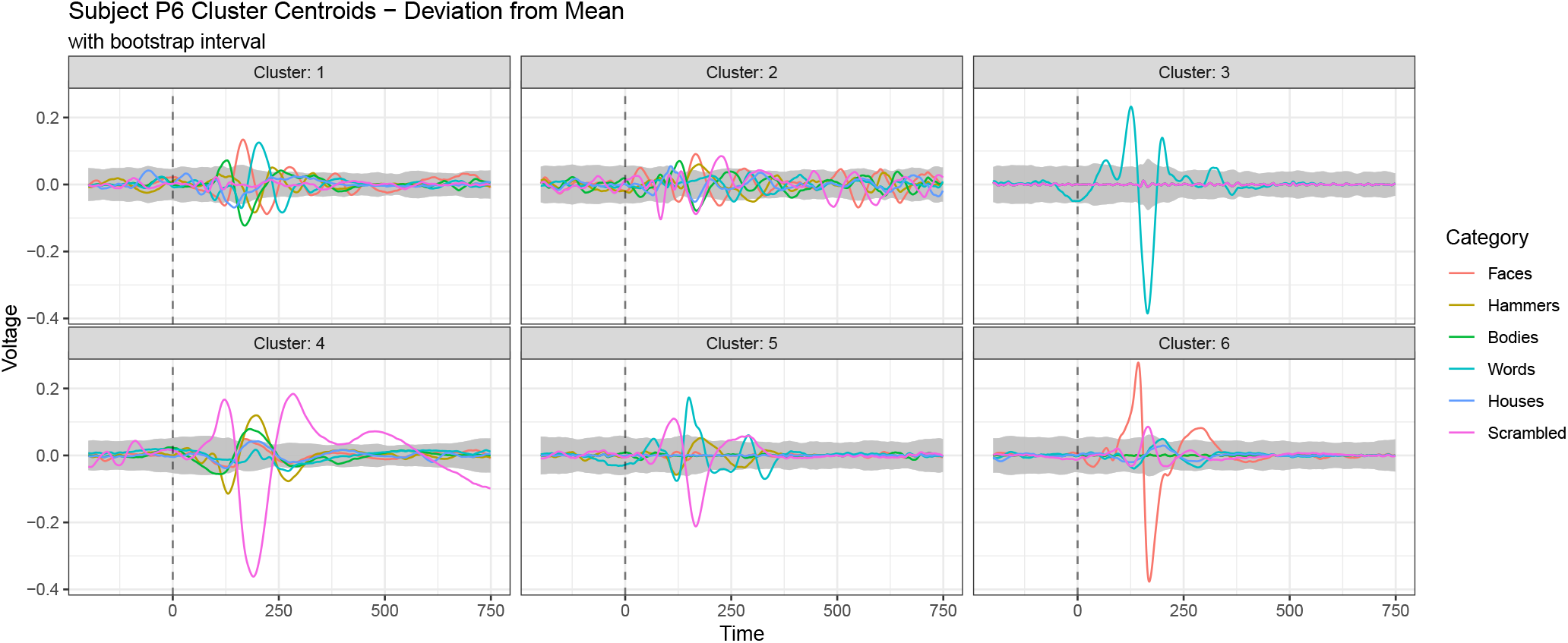
P6 Cluster Centroids: This plot shows the centroids of clusters with detected category sensitivity for subject P6. Dashed vertical lines indicate stimulus onset 500 ms into the trial recording. Gray bands in each plot indicate a time-varying 90% confidence region for the overall mean. Category sensitivity is assessed by comparing a category’s deviation from the mean at each time point to the width of the confidence band. The large deviations in clusters 1 and 6 suggest face sensitivity in the corresponding electrodes. Clusters 1, 3 and 5 show apparent word sensitivity, and clusters 4 and 5 show apparent sensitivity to scrambled faces.

### 3.3. Classification

For confirmation of the detected differences between categories in Section 3.2, we used the estimated centroids to attempt to classify a held out test set of trials.

#### 3.3.1. All-cluster Classification in iEEG

We first consider classifying test set trials using centroids estimated for all clusters. This method of classification is explained in Section 2.4. Accuracy was assessed using permutation tests described in Section 2.5.7. We compare the sensitivity and false positive rate for each category to the sensitivity, false positive rate trade-off corresponding to the 95^*th*^ percentile of the *d*′ permutation distribution. The results of the permutation tests for each subject and candidates for category sensitivity are shown in Figure 4. Aggregating across iEEG subjects, 21 of the 23 FWER-controlled candidates and 29 of the 32 FDR-controlled candidates for category sensitivity were corroborated by all-cluster classification success. The corroborated category sensitivity for P6 is shown in Figure 5.

**Figure 4:**
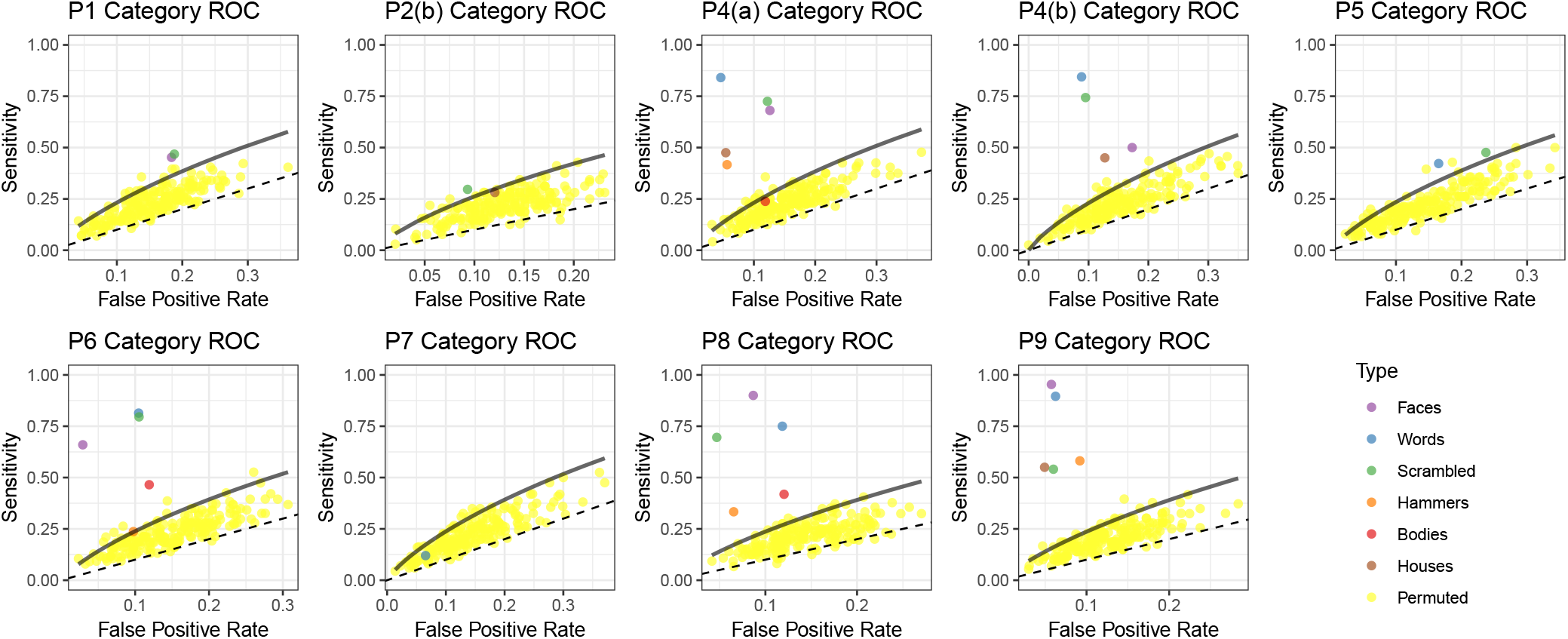
Permutation Test for All-Cluster Classification in iEEG: For each iEEG subject we evaluate six-way or seven-way classification success on the test set using a permutation test (Section 2.5.7). For each label shuffle, we find the best single-category *d*′. The permutation distribution is plotted along with the solid black line indicating the false positive rate, sensitivity trade-off corresponding to the 95^*th*^ percentile of the *d*′ permutation distribution. We evaluate classification results for categories selected by NSC with FDR control and see 29 of the 32 selected category-subject pairs out-perform the 95% cutoff for the corresponding subject.

**Figure 5:**
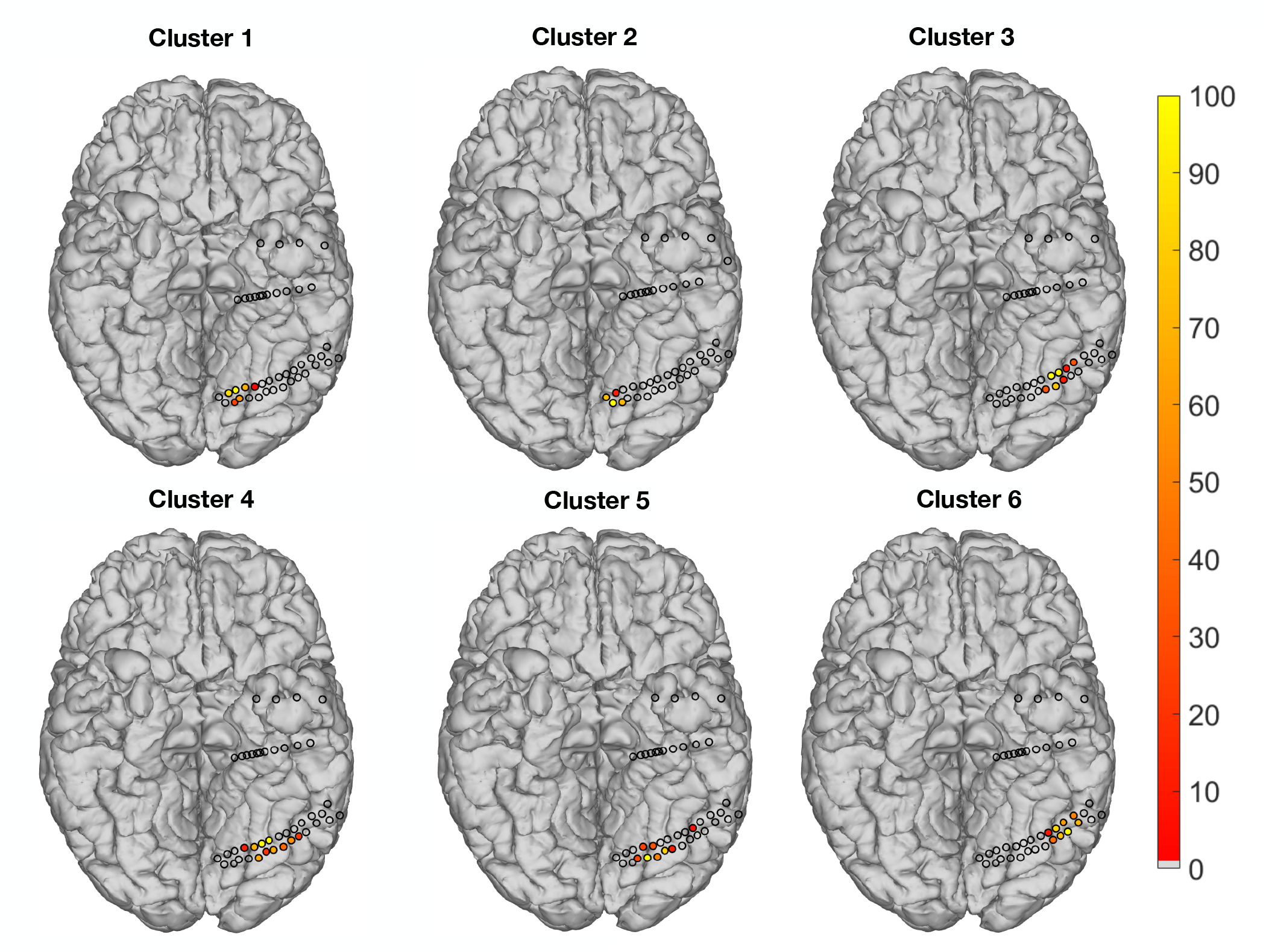
Anatomical locations of category-sensitive regions in P6: Circles indicate electrodes analyzed. The color of the circle indicates relative strength of membership in the given cluster, i.e. the magnitude of the coefficient in **W** (Section 2.2) relative to the highest magnitude coefficient for that cluster. Gray circles are electrodes that are not part of a category-sensitive cluster. As an example, cluster 6 is centered in the fusiform gyrus. Results from Figure 3 indicate face sensitivity in this area.

#### 3.3.2. Single-cluster Classification in iEEG

We also wish to attribute classification success to specific clusters. We repeated the evaluation in Section 3.3.1 without aggregating distance for classification across clusters, as described in Section 2.5.6. Each cluster then had its own classification score for each category. Table 3.3.2 shows for each iEEG subject the proportion of candidate category-cluster pairs that were corroborated by single-cluster classification and the categories for which there is corroborated sensitivity. Aggregating across iEEG subjects, we find that 36 of the 44 FWER-controlled candidate category-cluster pairs and 15 of the additional 31 FDR-controlled candidate category-cluster pairs were corroborated.

#### 3.3.3. Classification in MEG

We applied both all-cluster and single-cluster classification to three MEG subjects. Accuracy was assessed using permutation tests described in Section 2.5.7. The results of the permutation tests for each subject are shown in Figure 8. We find that 11 of the 12 categories across the three subjects have successful classification. Single-cluster classification found 27 cluster-category sensitive pairs in S1, 21 in S3 and 8 in S2. Figure 6 highlights several anatomical locations of category sensitivity in the three subjects, with corresponding centroids shown in Figure 7. These clusters correspond to spatially continuous regions ranging from 308 to 587 vertices.

**Figure 6:**
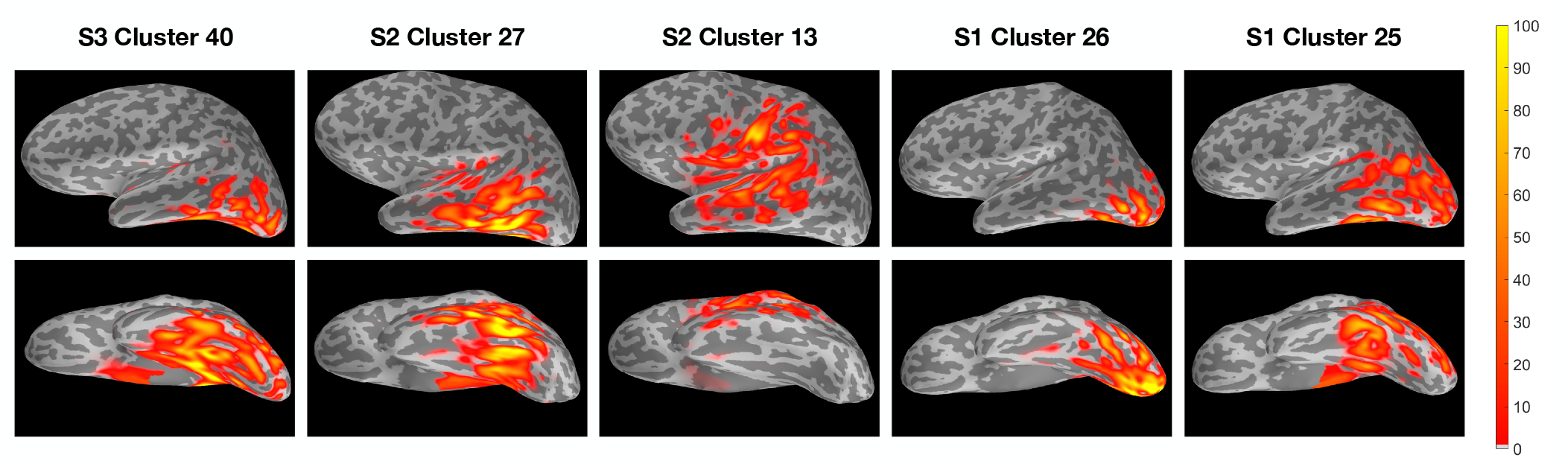
Anatomical locations of category-sensitive regions in MEG subjects: Each column shows an example category-sensitive region from lateral (top) and ventral (bottom) perspectives. Color indicates strength of cluster membership i.e. the magnitude of the coefficient in **W** (Section 2.2) relative to the highest magnitude coefficient in the cluster. As an example, cluster 27 in S2 is located in ventral temporal cortex and, from Figure 7, displays word sensitivity

**Figure 7:**
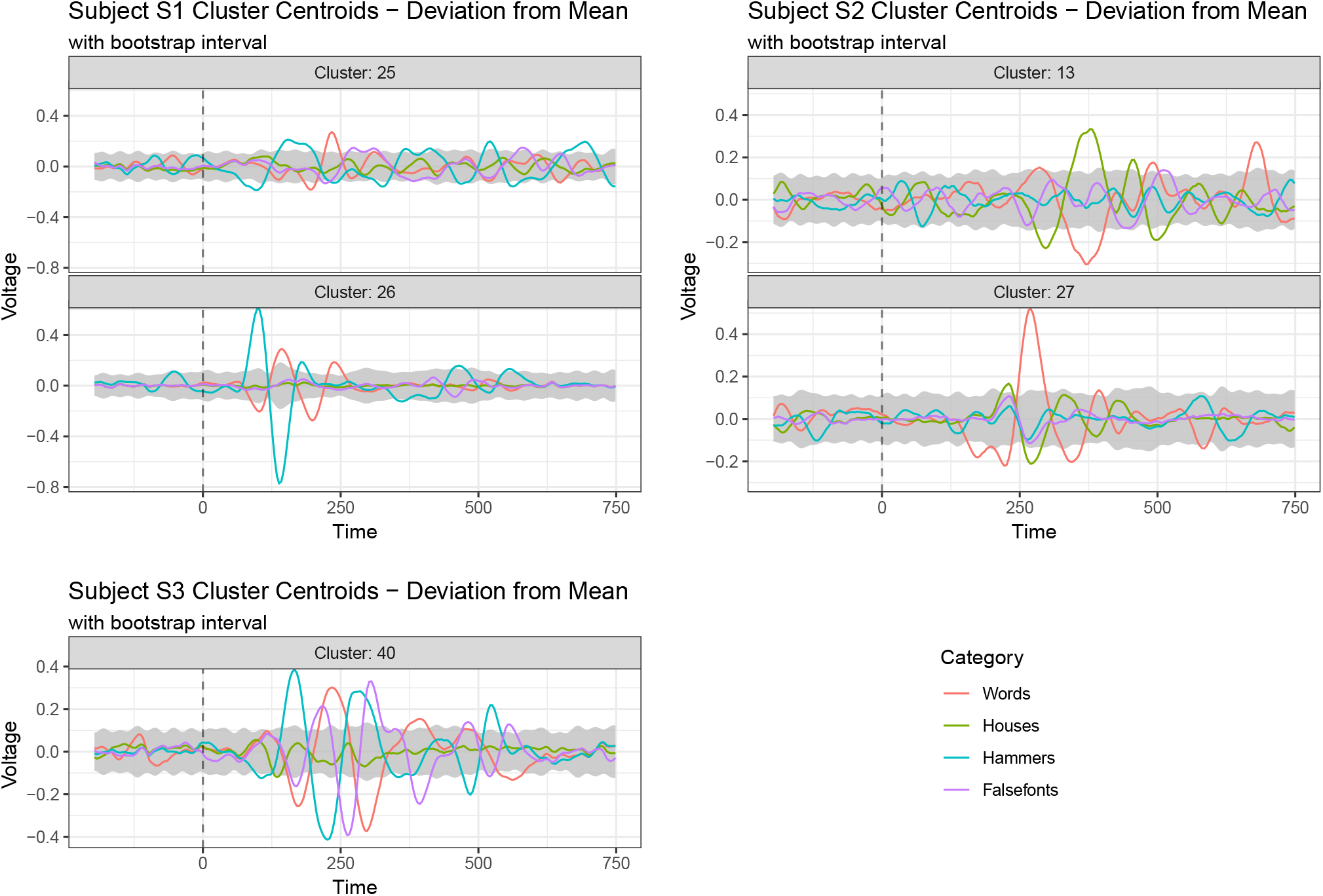
MEG Cluster Centroids: This plot shows the centroids of clusters with detected category sensitivity for MEG subjects. Dashed vertical lines indicate stimulus onset 500 ms into the trial recording. Gray bands in each plot indicate a time-varying 90% confidence region for the overall mean. Category sensitivity is assessed by comparing a category’s deviation from the mean at each time point to the width of the confidence band.

**Figure 8:**
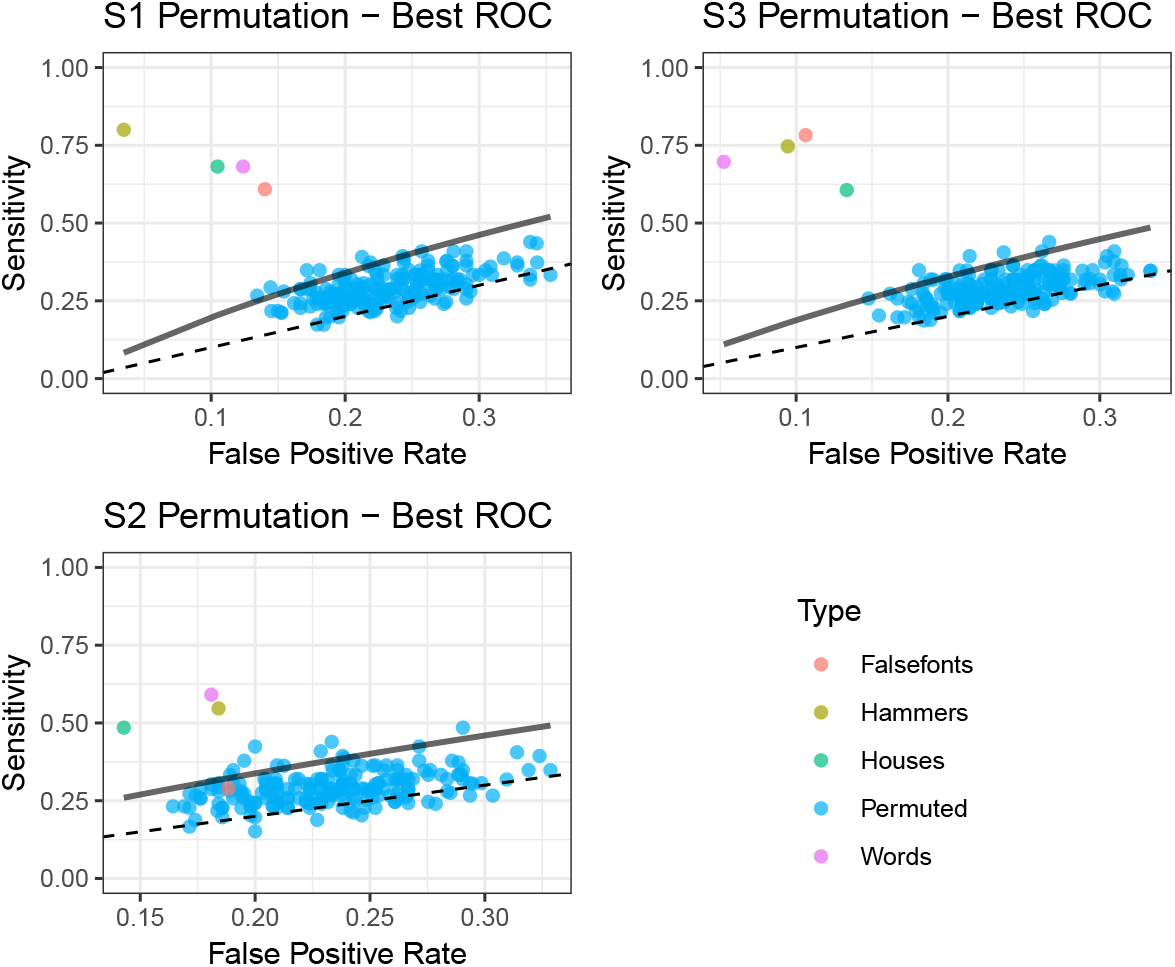
Permutation Test for All-Cluster Classification in MEG: For each MEG subject we evaluate four-way classification success on the test set using a permutation test (Section 2.5.7). For each label shuffle, we find the best single-category *d*′. The permutation distribution is plotted along with the solid black line indicating the false positive, sensitivity trade-off corresponding to the 95^*th*^ percentile of the permutation distribution. We find that 11 of the 12 category-subject pairs have successful classification.

S1 demonstrated two distinct clusters of word-sensitive vertices. The first peaked early, around 135 ms post-stimulus, and was concentrated around early visual cortices in the occipital lobe. The second cluster was slightly more anterior, in ventral temporal cortex and inferior temporal gyrus, and peaked at 240 ms. In S2, we also found two distinct clusters of word-sensitive vertices. One cluster concentrated primarily on ventral temporal cortex demonstrated relatively early activity for words, peaking at 275 ms. The other word-sensitive cluster was primarily concentrated on the left superior temporal sulcus, angular gyrus and ventral somatosensory cortex. The activity of these regions was later than that of the first cluster, peaking around 375 ms. Finally, S3 demonstrated a highly category-discriminant cluster of vertices encompassing several ventral visual stream areas including early visual cortex and ventral temporal cortex. The temporal dynamics of this cluster contained several oscillations of object discriminability for three of the four object categories.

#### 3.3.4. Results of Model Comparisons

Figure 9 compares classification in iEEG subjects across models (see Section 2.5.8). Our goal is to understand how tuning for interpretability, and its corresponding benefits for learning about underlying neural processing, affects predictive performance. We find that pre-stimulus based tuning shows comparable performance to using a prototype without dimensionality reduction and shrinkage, while cross validation provides the best predictive performance. Figure 10 compares classification in MEG subjects and finds simimilar predictive performance using SCOTS to classification without shrinkage after dimensionality reduction and classification using neither shrinkage nor dimensionality reduction.

**Figure 9:**
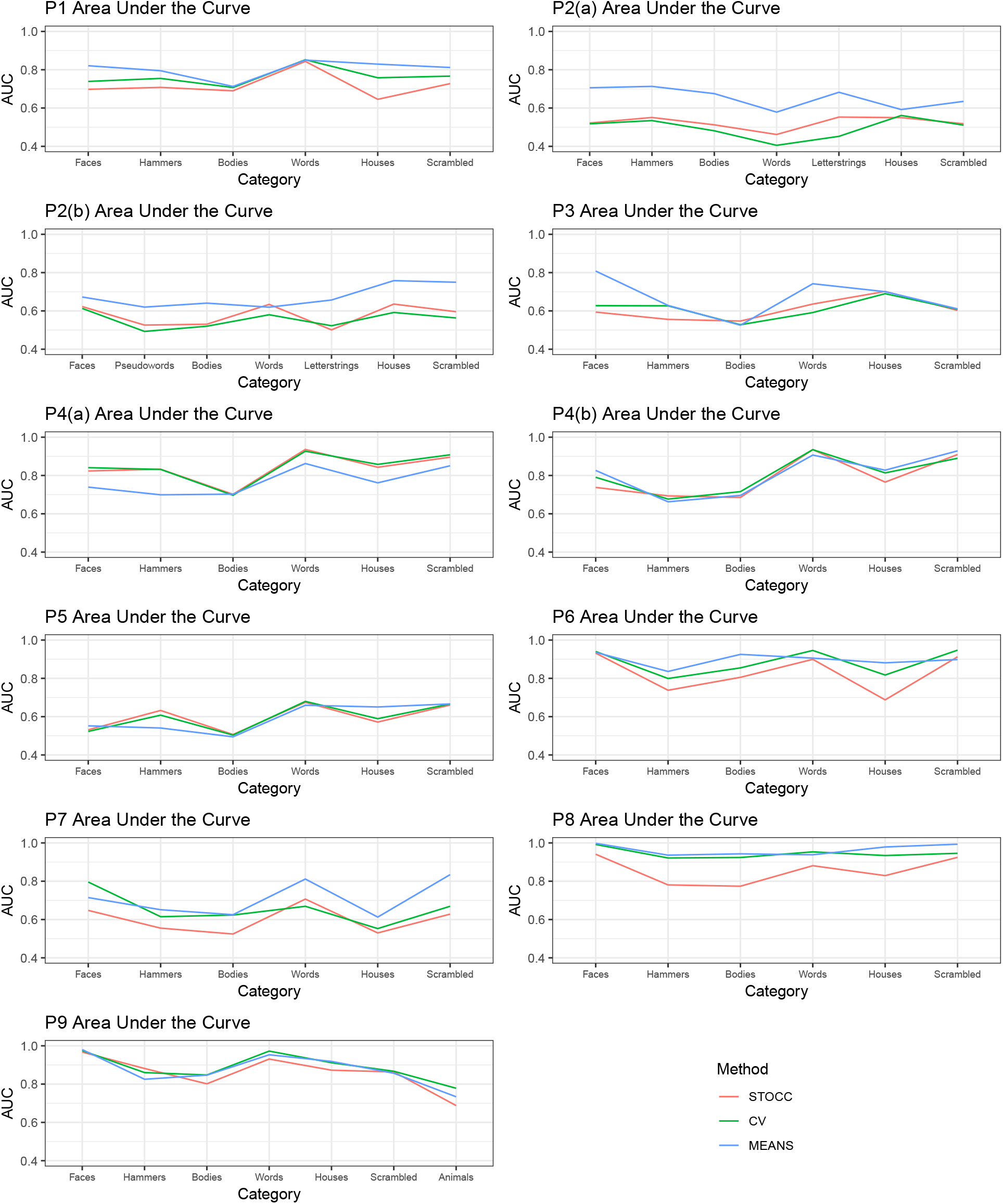
Category-wise AUC for Method Variants in iEEG: This figure compares category-wise ROC trade-offs in iEEG for SCOTS to a version with NSC penalty weights chosen by cross validation (CV) and a null form without either dimensionality reduction or regularization (MEANS). The results for SCOTS show comparable predictive performance even though it optimizes for interpretability.

**Figure 10:**
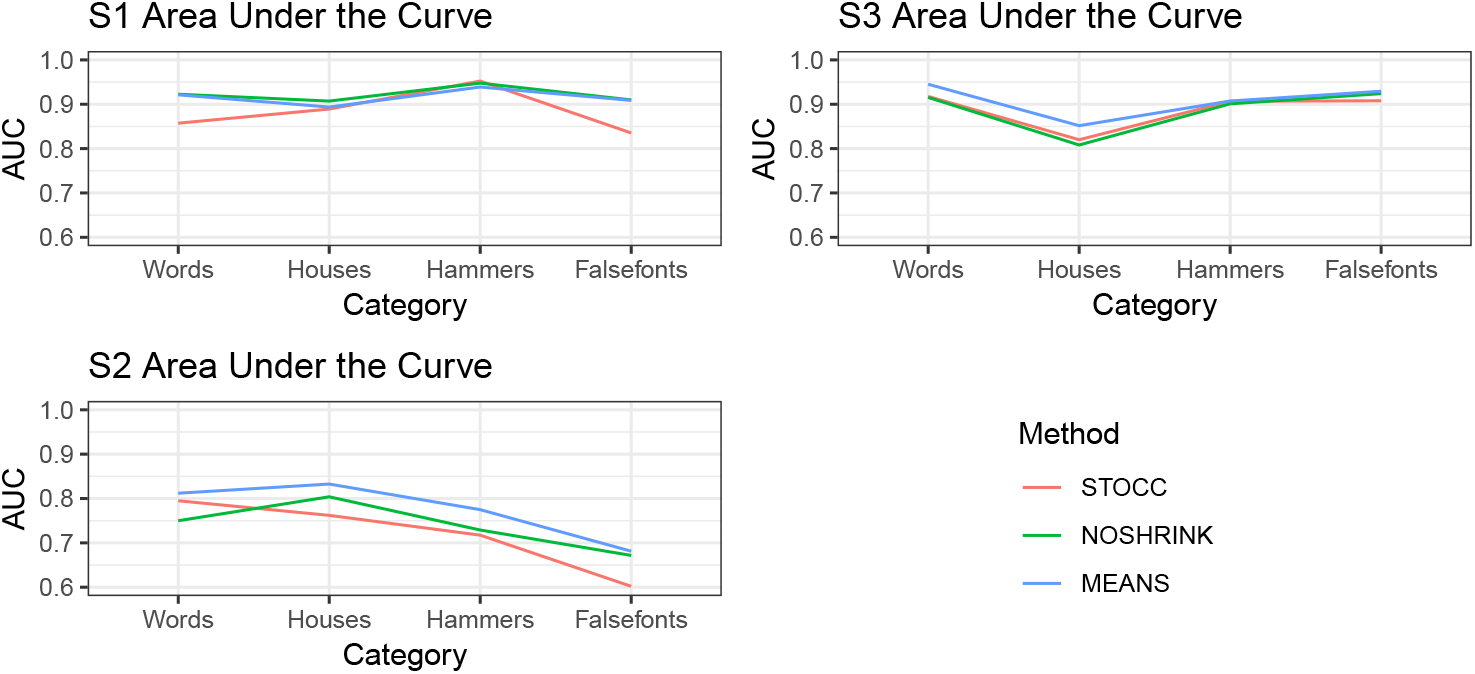
Category-wise AUC for Method Variants in MEG: We compare category-wise ROC trade-offs in MEG for SCOTS to a reduced form with only sparse SNMF (NOSHRINK) and a null form without either dimensionality reduction or regularization (MEANS). SCOTS optimizes for interpretability with little loss of predictive performance.

## 4. Discussion

One fundamental task in neuroscience is mapping brain function, both in terms of the specialization of different brain regions and their temporal latency. In this paper we proposed SCOTS, a method for finding neural activity specific to particular categories of visual stimuli that can be interpreted both spatially and temporally. Our model pairs two major components that are new to the problem of localizing neural activity: overlapping clustering as a means of data-driven aggregation of neural information and dimensionality reduction, and grouped nearest shrunken centroids in the wavelet domain for category sensitivity based cluster and time segment selection. These components are important for avoiding the problem of search inherent to methods currently in use. We also emphasize the use of wavelets to adjust both the sparse SNMF and grouped NSC to time series.

Our results show that the proposed method is successful, with eight cases of category sensitivity validated across six iEEG subjects and 14 cases of clusters with validated category sensitivity. Subjects P6, P1, P2 and P3 all demonstrated face sensitivity in the fusiform gyrus, which has previously been implicated in processing face forms (Kanwisher et al., 1997). P4(b) demonstrates face sensitivity in the right homologue of this region. P6 also demonstrated word sensitivity in the left fusiform gyrus, consistent with previous fMRI work on word processing in the ventral stream (McCandliss et al., 2003).

In MEG subjects, the temporal dynamics of their category-discriminant responses often varied as expected with their anatomical location. S1 and S3 both displayed evidence of hierarchical visual processing (Grill-Spector and Weiner, 2014): early visual clusters respond earlier than ventral temporal clusters in S1, and in S3 a highly category-discriminant cluster of vertices across several ventral visual stream areas includes early visual cortex and ventral temporal cortex. In S2, early activity in the ventral temporal cortex in response to words was also consistent with previously noted importance of regions within ventral temporal cortex to visual word processing (McCandliss et al., 2003). Later activity concentrated on the left superior temporal sulcus, angular gyrus and ventral somatosensory cortex may reflect post-lexical processing of word stimuli (Fiez and Petersen, 1998).

Sparse SNMF and NSC are also related to other methods of multivariate analysis. Our adjustment to NMF can be seen as a structured form of principal component analysis (PCA) - in fact the SVD-based initialization we use is equivalent to PCA. Our use of sparsity in SNMF allows us to better interpret components as locally co-occurring neural activity across recording channels. While PCA describes statistically independent modes of variation in the data, our use of sparse SNMF does not require statistical independence. This leaves open the question of how these separate components interact with each other. Further investigation could reveal dynamic interactions between different brain regions that contribute to the processing of different object categories.

Nearest shrunken centroids is also a structured version of other popular classification methods. When Euclidean distance is used and centroids are not shrunken, this incorporates assumptions from both naïve Bayes and linear discriminant analysis (LDA), i.e. observations are normally distributed with covariance identical between classes and all features mutually independent. While this is a highly unreasonable assumption for observations in a time series, our wavelet transformation shifts the implicit independence assumption to the wavelet domain. A natural avenue for future investigation would be to weaken these assumptions for greater flexibility, though this may also require experiments with greater numbers of trials.

It is also important to note that this approach examines only category sensitivity found in the time domain. Similar methods could be used to find information carried in the frequency domain. This oscillatory behavior can be characterized using a short-time Fourier transform so that each trial is associated with a nonnegative tensor indexed by time, frequency and recording channel. Adapting SCOTS to this data could reveal regions in time-frequency space with class-specific behavior or find class-specific spectral signatures that emerge at varying temporal latencies.

In this paper we demonstrate the application of SCOTS to iEEG and MEG data, and in principle similar methodology could be applied for other human electrophysiological data, including scalp EEG. One of the primary differences between iEEG and scalp EEG/MEG is the spatial configuration of recording locations – both scalp EEG and MEG offer whole brain coverage with more uniform coverage density compared to iEEG. As such, the relatively sharp edges in clusters induced by the lasso penalty may be unrealistic. To instead get tapered cluster edges, using an elastic net penalty (combining both *L*_1_ and *L*_2_ terms) may be preferred. Another option is a graph-based penalty, encouraging similarity in loadings based on spatial distance.

We also advocate more generally for dimensionality reduction based analysis techniques as in Cunningham and Yu (2014). These techniques provide a natural domain for analyzing dynamics across a recorded system. In our case, overlapping clustering gives us a way to handle shared behavior across recording channels and separate distinct sources within recording channels where necessary. The clusters we get are then more relevant to distinguishing responses to stimuli.

Clusters or factors from dimensionality reduction should also be useful for other tasks. One challenge with intracranial EEG is the idiosyncratic placement of electrodes across subjects, which makes it difficult to compare results across subjects. Clusters or factors derived from dimensionality reduction could serve as a way to bridge this gap between subjects. For example, one could find clusters of neural activity that behave similarly across subjects. If these clusters are localized to nearby regions of space across subjects, they may be thought of as equivalent, offering a way to pool information in separate subjects. Further, defining equivalent areas in cluster space would likely be more robust than using individual electrodes.

## Appendix A. Optimization Procedure

### Appendix A.1. SNMF

We use alternating minimization, summarized in Algorithm 1, to reach a local minimum of (1), which is a non-convex minimization problem. We alternate between updating estimates for **W** and **F**_1_,…, **F**_*n*_. Updating each **F**_*i*_ requires solving a series of ridge regressions in constrained form. This can be done using the Frank-Wolfe method. The update for **W** is a nonnegative lasso problem. This can be solved using proximal gradient descent (Hastie et al., 2015) (also known as iterative shrinkage-thresholding (Beck and Teboulle, 2009)), with steps given by the proximal function

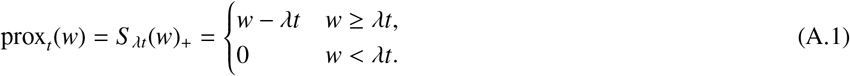

#### Algorithm 1 Sparse Semi-NMF with Wavelets

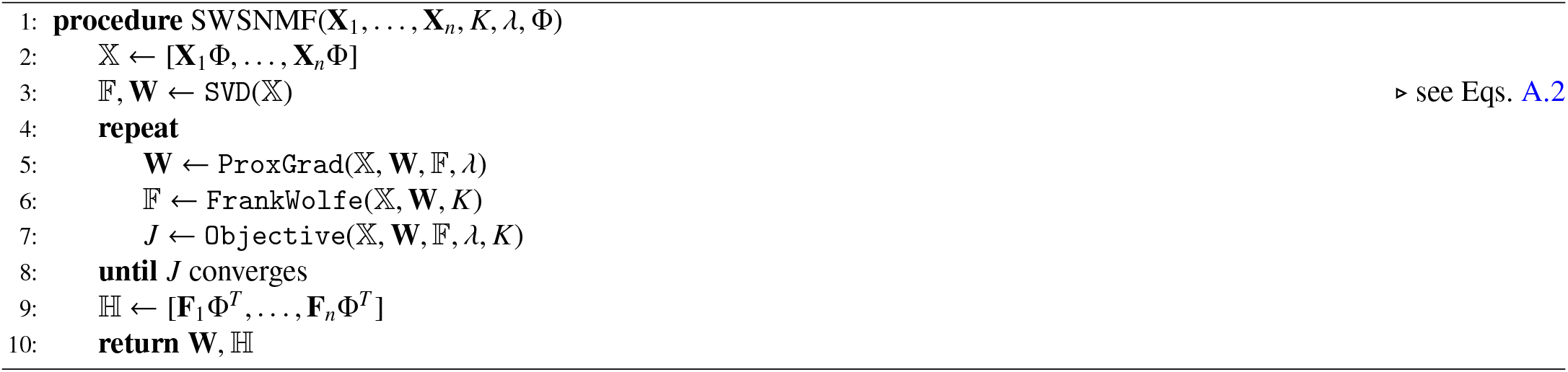

We initialize estimates using the singular value decomposition (SVD). We first horizontally concatenate trials, writing 𝕏 = [**X**_1_Φ,…, **X**_*n*_Φ]. Then using **A**_[*ℓ,m*]_ to denote the first *ℓ* rows and *m* columns of a matrix **A**, we set:

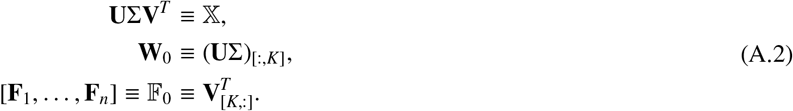

Note that column signs of **U** and **V** in the SVD can be arbitrary, and we choose signs so that the column sums of **W**_0_ are each positive, i.e. to maximize cluster weights after setting negative values to 0.

### Appendix A.2. WNSC

This is solved using alternating minimization in each **F**^*g*^ and 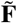, as summarized in Algorithm 2. We perform minimization in **F**^*g*^ using the **R** `SGL` package (Simon et al., 2013). The minimizer in 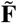 is a pointwise median across category centroids.

## Appendix B. Outlier Removal

Outlying trials were detected using a two-step procedure. First, for each trial the average time signal across all channels was calculated. Second, for each trial the maximum absolute signal across time was found and *z*-scored with respect to the distribution of such maximums across trials. All trials with a *z*-score above 4 were removed.

### Algorithm 2 Wavelet Nearest Shrunken Centroids

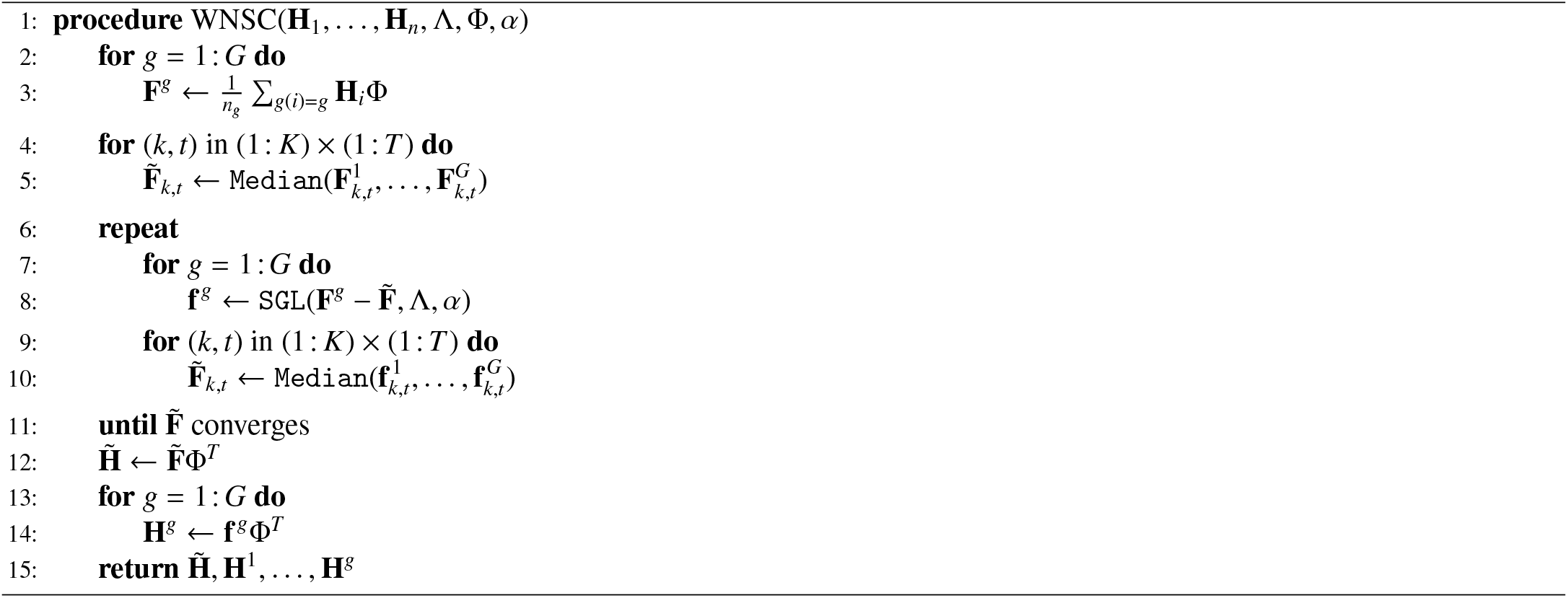

## Appendix C. Additional Results

A representative choice of figures is included in the results. Here we present all cluster centroids and corresponding cluster locations with NSC-selected sensitivity verified by out of sample classification.

### Appendix C.1. iEEG Centroids

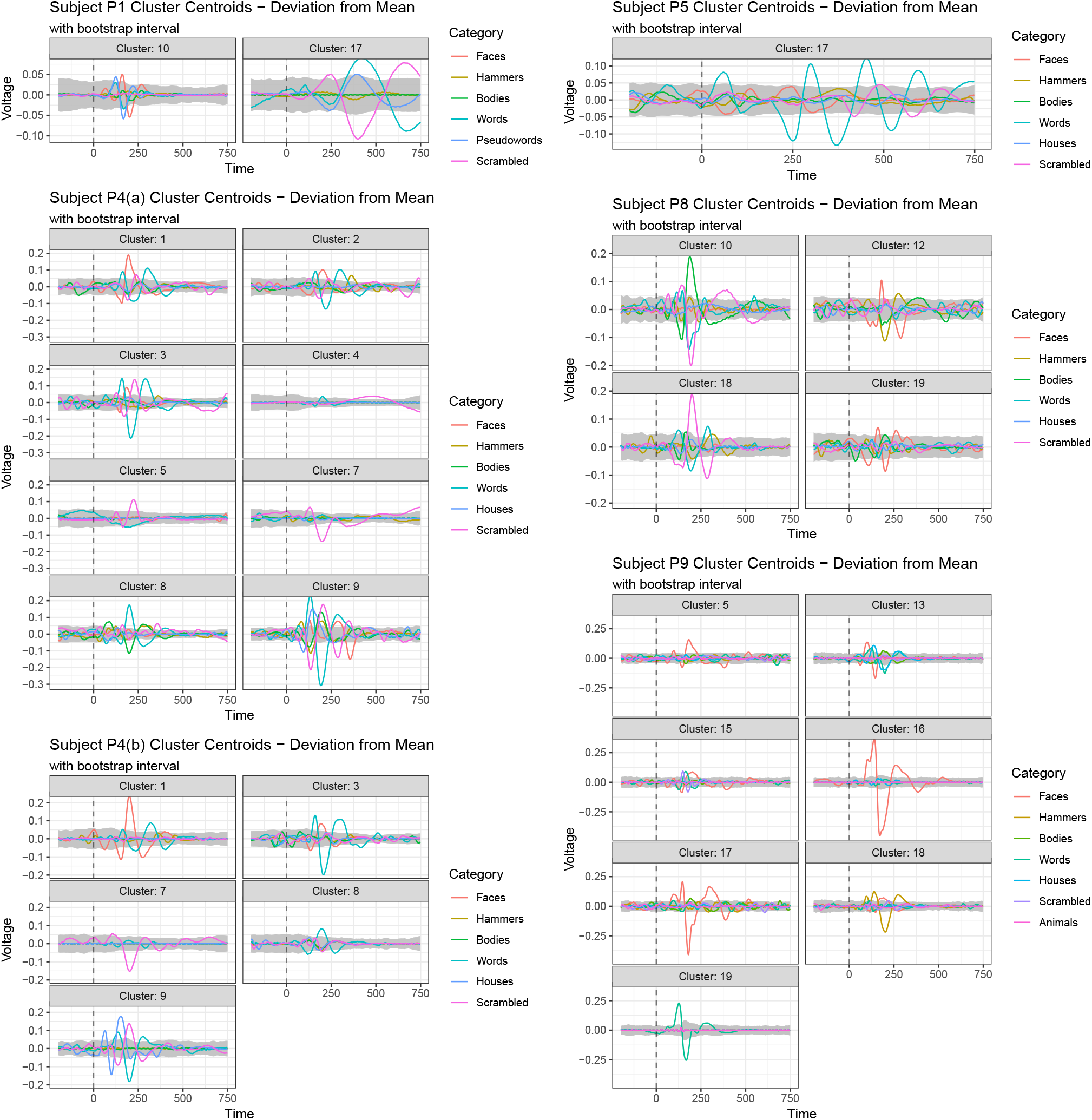

### Appendix C.2. iEEG Cluster Locations

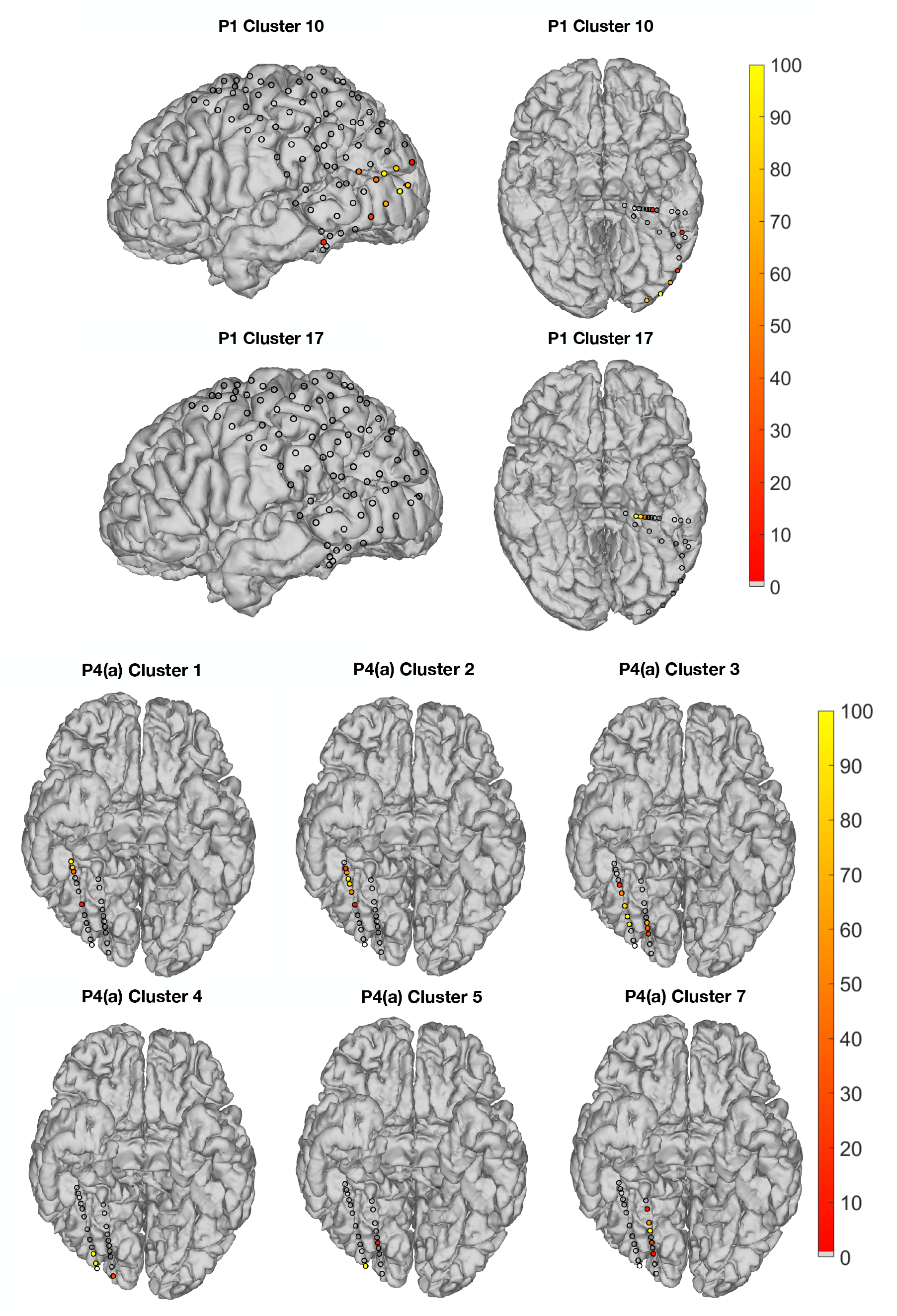
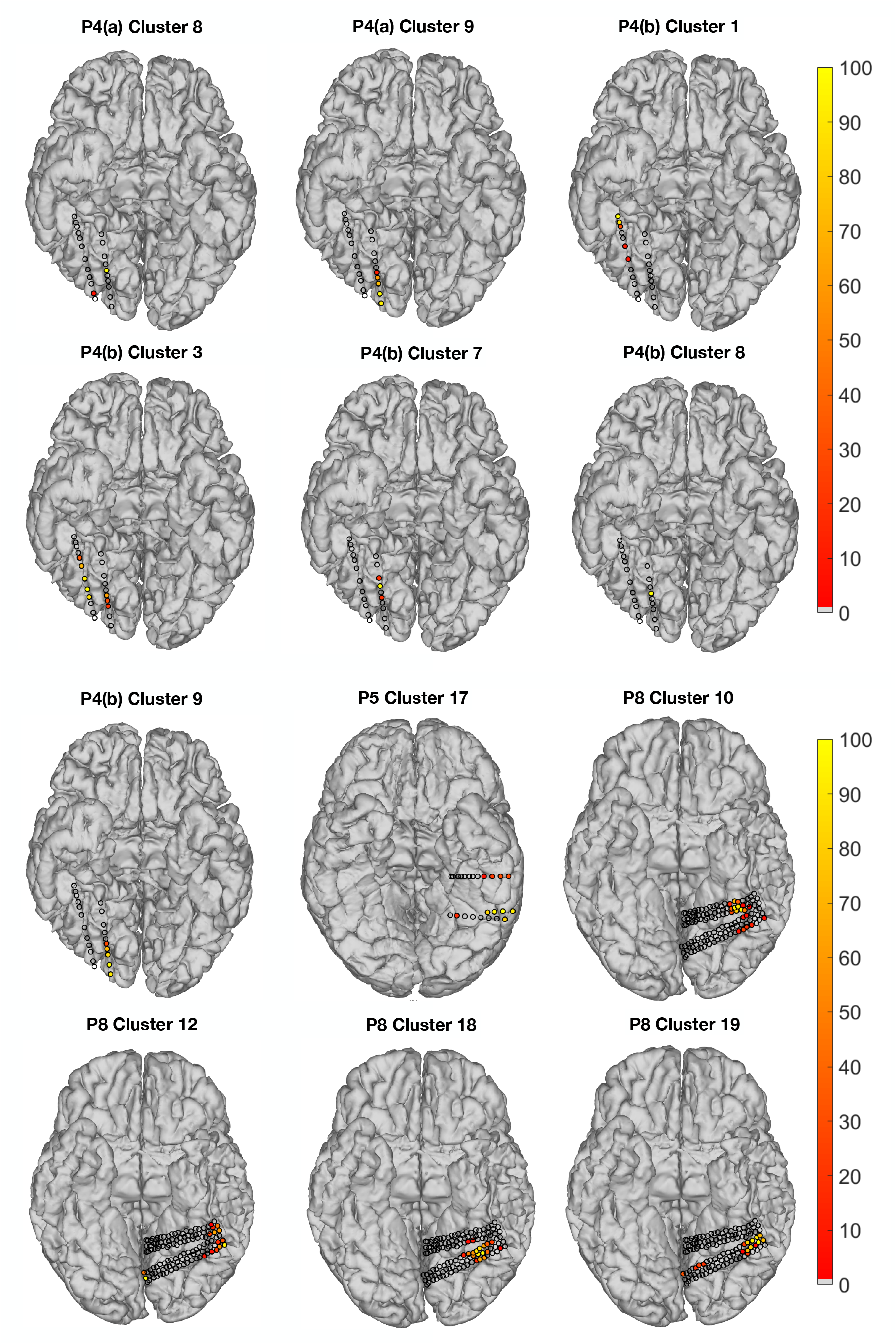
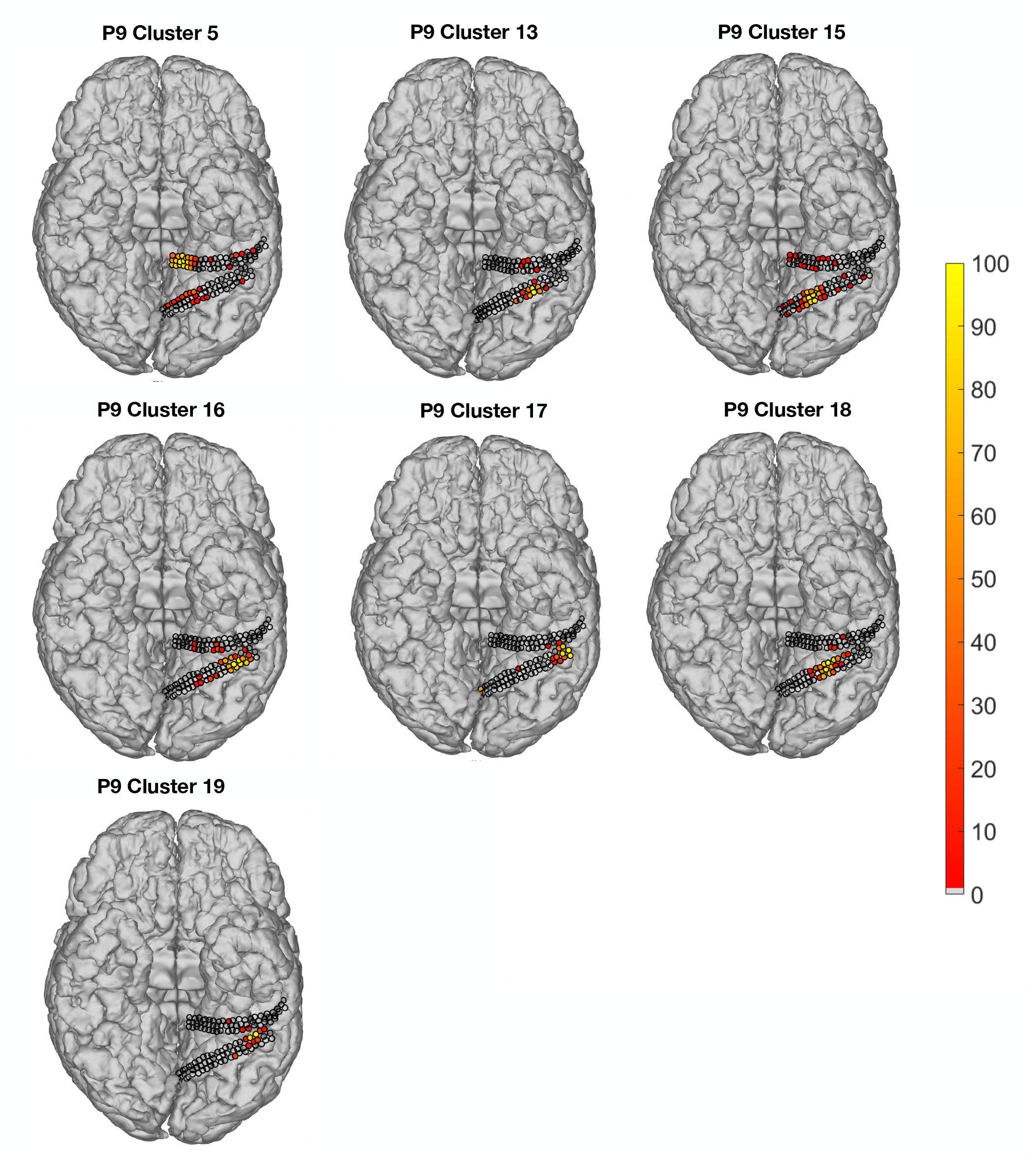

## References

Allison, T., Ginter, H., McCarthy, G., Nobre, A. C., Puce, A., Luby, M., Spencer, D. D., 1994. Face recognition in human extrastriate cortex. J Neurophysiol 71 (2), 821–825.

Allison, T., Puce, A., Spencer, D. D., McCarthy, G., 1999. Electrophysiological studies of human face perception. i: Potentials generated in occipitotemporal cortex by face and non-face stimuli. Cereb Cortex 9 (5), 415–430.

Beck, A., Teboulle, M., 2009. A fast iterative shrinkage-thresholding algorithm for linear inverse problems. SIAM J Imaging Sci 2 (1), 183–202.

Benjamini, Y., Hochberg, Y., 1995. Controlling the false discovery rate: a practical and powerful approach to multiple testing. J R Stat Soc Series B Stat Methodol 57 (1), 289–300.

Buzsáki, G., Anastassiou, C. A., Koch, C., 2012. The origin of extracellular fields and currents - eeg, ecog, lfp and spikes. Nat Rev Neurosci 13 (6), 407.

Cichocki, A., Phan, A. H., Zdunek, R., Zhang, L.-Q., 2007. Flexible component analysis for sparse, smooth, nonnegative coding or representation. In: International Conference on Neural Information Processing. Springer, pp. 811–820.

Cunningham, J. P., Yu, B. M., 2014. Dimensionality reduction for large-scale neural recordings. Nat Neurosci 17 (11), 1500.

Devarajan, K., 2008. Nonnegative matrix factorization: an analytical and interpretive tool in computational biology. PLoS Comput Biol 4 (7), e1000029.

Ding, C. H., Li, T., Jordan, M. I., 2010. Convex and semi-nonnegative matrix factorizations. IEEE Trans Pattern Anal Mach Intell 32 (1), 45–55.

Donoho, D. L., Johnstone, J. M., 1994. Ideal spatial adaptation by wavelet shrinkage. Biometrika 81 (3), 425–455.

Drakakis, K., Rickard, S., De Fréin, R., Cichocki, A., 2008. Analysis of Financial Data Using Non-Negative Matrix Factorization. International Mathematical Forum 3 (38), 1853–1870.

Etzel, J. A., Zacks, J. M., Braver, T. S., 2013. Searchlight analysis: promise, pitfalls, and potential. Neuroimage 78, 261–269.

Févotte, C., Bertin, N., Durrieu, J.-L., 2009. Nonnegative matrix factorization with the itakura-saito divergence: With application to music analysis. Neural Comput 21 (3), 793–830.

Fiez, J. A., Petersen, S. E., 1998. Neuroimaging studies of word reading. Proc Natl Acad Sci USA 95 (3), 914–921.

Ghuman, A. S., Brunet, N. M., Li, Y., Konecky, R. O., Pyles, J. A., Walls, S. A., Destefino, V., Wang, W., Richardson, R. M., 2014. Dynamic encoding of face information in the human fusiform gyrus. Nat Commun 5, 5672.

Gramfort, A., Luessi, M., Larson, E., Engemann, D. A., Strohmeier, D., Brodbeck, C., Parkkonen, L., Hämäläinen, M. S., 2014. Mne software for processing meg and eeg data. Neuroimage 86, 446–460.

Grill-Spector, K., Weiner, K. S., 2014. The functional architecture of the ventral temporal cortex and its role in categorization. Nat Rev Neurosci 15 (8), 536.

Hamilton, L. S., Edwards, E., Chang, E. F., 2018. A spatial map of onset and sustained responses to speech in the human superior temporal gyrus. Curr Biol 28 (12), 1860–1871.

Hastie, T., Tibshirani, R., Wainwright, M., 2015. Statistical Learning with Sparsity: the Lasso and Generalizations. CRC Press.

Haufe, S., Meinecke, F., Görgen, K., Dähne, S., Haynes, J.-D., Blankertz, B., Bießmann, F., 2014. On the interpretation of weight vectors of linear models in multivariate neuroimaging. Neuroimage 87, 96–110.

Hermes, D., Miller, K. J., Noordmans, H. J., Vansteensel, M. J., Ramsey, N. F., 2010. Automated electrocorticographic electrode localization on individually rendered brain surfaces. J Neurosci Methods 185 (2), 293–298.

Hirshorn, E. A., Li, Y., Ward, M. J., Richardson, R. M., Fiez, J. A., Ghuman, A. S., 2016. Decoding and disrupting left midfusiform gyrus activity during word reading. Proc Natl Acad Sci USA 113 (29), 8162–8167.

Kanwisher, N., McDermott, J., Chun, M. M., 1997. The fusiform face area: a module in human extrastriate cortex specialized for face perception. J Neurosci 17 (11), 4302–4311.

Kravitz, D. J., Saleem, K. S., Baker, C. I., Ungerleider, L. G., Mishkin, M., 2013. The ventral visual pathway: an expanded neural framework for the processing of object quality. Trends Cogn Sci 17 (1), 26–49.

Kriegeskorte, N., Douglas, P. K., 2018. Cognitive computational neuroscience. Nat Neurosci, 1.

Kriegeskorte, N., Goebel, R., Bandettini, P., 2006. Information-based functional brain mapping. Proc Natl Acad Sci USA 103 (10), 3863–3868.

Lee, D. D., Seung, H. S., 1999. Learning the parts of objects by non-negative matrix factorization. Nature 401 (6755), 788.

Li, Y., Richardson, R. M., Ghuman, A. S., 2017. Multi-connection pattern analysis: Decoding the representational content of neural communication. Neuroimage 162, 32–44.

Liu, H., Agam, Y., Madsen, J. R., Kreiman, G., 2009. Timing, timing, timing: fast decoding of object information from intracranial field potentials in human visual cortex. Neuron 62 (2), 281–290.

Macmillan, N. A., Creelman, C. D., 2004. Detection Theory: A User’s Guide. Psychology Press.

McCandliss, B. D., Cohen, L., Dehaene, S., 2003. The visual word form area: expertise for reading in the fusiform gyrus. Trends Cogn Sci 7 (7), 293–299.

McCarthy, G., Puce, A., Belger, A., Allison, T., 1999. Electrophysiological studies of human face perception. ii: Response properties of face-specific potentials generated in occipitotemporal cortex. Cereb Cortex 9 (5), 431–444.

Naselaris, T., Kay, K. N., Nishimoto, S., Gallant, J. L., 2011. Encoding and decoding in fmri. Neuroimage 56 (2), 400–410.

Nobre, A. C., Allison, T., McCarthy, G., 1994. Word recognition in the human inferior temporal lobe. Nature 372 (6503), 260.

Oostenveld, R., Fries, P., Maris, E., Schoffelen, J.-M., 2011. Fieldtrip: open source software for advanced analysis of meg, eeg, and invasive electrophysiological data. Comput Intell Neurosci 2011, 1.

Ortega-Martorell, S., Lisboa, P. J., Vellido, A., Simões, R. V., Pumarola, M., Julià-Sapé, M., Arús, C., 2012. Convex non-negative matrix factorization for brain tumor delimitation from mrsi data. PLoS One 7 (10), e47824.

Rasmussen, P. M., Hansen, L. K., Madsen, K. H., Churchill, N. W., Strother, S. C., 2012. Model sparsity and brain pattern interpretation of classification models in neuroimaging. Pattern Recognit 45 (6), 2085–2100.

Sajda, P., Du, S., Brown, T. R., Stoyanova, R., Shungu, D. C., Mao, X., Parra, L. C., 2004. Nonnegative matrix factorization for rapid recovery of constituent spectra in magnetic resonance chemical shift imaging of the brain. IEEE Trans Med Imaging 23 (12), 1453–1465.

Simon, N., Friedman, J., Hastie, T., Tibshirani, R., 2013. SGL: Fit a GLM (or Cox model) with a Combination of Lasso and Group Lasso Regularization. R package version 1.1. URL https://CRAN.R-project.org/package=SGL

Su, L., Fonteneau, E., Marslen-Wilson, W., Kriegeskorte, N., 2012. Spatiotemporal searchlight representational similarity analysis in emeg source space. In: Int Workshop Pattern Recognit Neuroimaging. IEEE, pp. 97–100.

Tadel, F., Baillet, S., Mosher, J. C., Pantazis, D., Leahy, R. M., 2011. Brainstorm: a user-friendly application for meg/eeg analysis. Comput Intell Neurosci 2011, 8.

Taulu, S., Hari, R., 2009. Removal of magnetoencephalographic artifacts with temporal signal-space separation: demonstration with single-trial auditory-evoked responses. Hum Brain Mapp 30 (5), 1524–1534.

Taulu, S., Simola, J., 2006. Spatiotemporal signal space separation method for rejecting nearby interference in meg measurements. Phys Med Biol 51 (7), 1759.

Tesche, C., Uusitalo, M., Ilmoniemi, R., Huotilainen, M., Kajola, M., Salonen, O., 1995. Signal-space projections of meg data characterize both distributed and well-localized neuronal sources. Electroencephalogr Clin Neurophysiol 95 (3), 189–200.

Tibshirani, R., Hastie, T., Narasimhan, B., Chu, G., 2003. Class prediction by nearest shrunken centroids, with applications to dna microarrays. Stat Sci, 104–117.

Uusitalo, M. A., Ilmoniemi, R. J., 1997. Signal-space projection method for separating meg or eeg into components. Med Biol Eng Comput 35 (2), 135–140.

Virtanen, T., 2007. Monaural sound source separation by nonnegative matrix factorization with temporal continuity and sparseness criteria. IEEE Trans Audio Speech Lang Process 15 (3), 1066–1074.

Wang, F., Li, T., Wang, X., Zhu, S., Ding, C., 2011. Community discovery using nonnegative matrix factorization. Data Min Knowl Discov 22 (3), 493–521.

Wang, Y.-X., Zhang, Y.-J., 2013. Nonnegative matrix factorization: A comprehensive review. IEEE Trans Knowl Data Eng 25 (6), 1336–1353.

Xu, W., Liu, X., Gong, Y., 2003. Document clustering based on non-negative matrix factorization. In: Proceedings of the 26th Annual International ACM SIGIR Conference on Research and Development in Information Retrieval. ACM, pp. 267–273.

Yamashita, O., Sato, M.-A., Yoshioka, T., Tong, F., Kamitani, Y., 2008. Sparse estimation automatically selects voxels relevant for the decoding of fmri activity patterns. Neuroimage 42 (4), 1414–1429.

Yokota, T., Zdunek, R., Cichocki, A., Yamashita, Y., 2015. Smooth nonnegative matrix and tensor factorizations for robust multi-way data analysis. Signal Processing 113, 234–249.

Zhao, Y., Ogden, R. T., Reiss, P. T., 2012. Wavelet-based lasso in functional linear regression. J Comput Graph Stat 21 (3), 600–617.

